# Single Cell Profiling Distinguishes Leukemia-Selective Chemotypes

**DOI:** 10.1101/2024.05.01.591362

**Authors:** Hannah L. Thirman, Madeline J. Hayes, Lauren E. Brown, John A. Porco, Jonathan M. Irish

## Abstract

A central challenge in chemical biology is to distinguish molecular families in which small structural changes trigger large changes in cell biology. Such families might be ideal scaffolds for developing cell-selective chemical effectors – for example, molecules that activate DNA damage responses in malignant cells while sparing healthy cells. Across closely related structural variants, subtle structural changes have the potential to result in contrasting bioactivity patterns across different cell types. Here, we tested a 600-compound Diversity Set of screening molecules from the Boston University Center for Molecular Discovery (BU-CMD) in a novel phospho-flow assay that tracked fundamental cell biological processes, including DNA damage response, apoptosis, M-phase cell cycle, and protein synthesis in MV411 leukemia cells. Among the chemotypes screened, synthetic congeners of the rocaglate family were especially bioactive. In follow-up studies, 37 rocaglates were selected and deeply characterized using 12 million additional cellular measurements across MV411 leukemia cells and healthy peripheral blood mononuclear cells. Of the selected rocaglates, 92% displayed significant bioactivity in human cells, and 65% selectively induced DNA damage responses in leukemia and not healthy human blood cells. Furthermore, the signaling and cell-type selectivity were connected to structural features of rocaglate subfamilies. In particular, three rocaglates from the rocaglate pyrimidinone (RP) structural subclass were the only molecules that activated exceptional DNA damage responses in leukemia cells without activating a detectable DNA damage response in healthy cells. These results indicate that the RP subset should be extensively characterized for anticancer therapeutic potential as it relates to the DNA damage response. This single cell profiling approach advances a chemical biology platform to dissect how systematic variations in chemical structure can profoundly and differentially impact basic functions of healthy and diseased cells.

## Introduction

Improving the quality of per-cell biological measurements has been a focus of cytometry over the last 20 years and has led to significant improvements in understanding cellular processes governing life, death, and specification of cell identity ^1–4^. However, in preclinical drug discovery and chemical biology it is common to use either well-based readouts, or alternatively, to deploy imaging or label free approaches that are neither single cell nor multiplexed, or lack in resolution or throughput. Suspension cytometry assays could potentially address these issues by balancing these factors while quantifying multiple features per cell ^5,6^.

Phospho-specific flow cytometry (phospho-flow) was developed to quantify a range of cellular functions at the single cell level both within and across different populations of cells in parallel ^7–9^. The technique involves quantifying one or more features of individual cells that have been stained with a panel of fluorescently tagged antibodies directed against specific epitopes, such as surface proteins, transcription factors, and phospho-proteins. This approach can therefore assess multiple signaling pathways simultaneously in mixtures of cells without the need to physically isolate different cell subsets present in human tissue samples. Thus, phospho-flow offers a platform for chemical biology that combines clinical relevance and multi-parameter single cell data. Throughput, consistency, and costs have been improved by a technique called fluorescent cell barcoding (FCB) that labels cells from a given well or sample with a unique signature of fluorescent dye levels - a barcode - so that they can be mixed, stained, and analyzed as a single sample ^10–12^. Phospho-flow and FCB protocols have evolved over the last two decades to closely align with drug discovery and translational research workflows ^13–15^. Phospho-flow-based single cell profiling has been used to screen sets of small molecules and metabolite extracts and has identified pathway- and cell-selective inhibitors in cell lines and primary samples ^3,6,16–19^. These prior studies focused on sets of diverse molecules; thus, there is an opportunity to explore multiplexed phospho-flow as a platform for dissecting structure-activity relationships (SAR).

Here, we aimed to develop bench and computational methods to apply single cell profiling to SAR studies. This approach aims to draw on quantitative structure-activity relationship (QSAR) approaches ^20^ and bring in single cell bench techniques and modern data science tools that draw on machine learning. Computational methods like t-distributed stochastic neighbor embedding (t-SNE) ^21–23^, UMAP ^24,25^, and PHATE ^26^ are used to analyze cytometry data ^24^ and are now growing in use in computational chemical biology ^6,16,26^. Recently developed algorithms, such as Tracking Responders EXpanding (T-REX) and Marker Enrichment Modeling (MEM), might be used to augment analysis and facilitate identification of rare cells and quantification of contextual protein enrichment ^27,28^. Tools like these that can resolve cellular heterogeneity and consider many readouts simultaneously may help to reveal connections between chemical structure and cellular bioactivity and accelerate development of molecules that selectively modulate chosen cell subsets and improve patient outcomes.

Here, we describe a multiplexed, single cell phospho-flow-based test of a chemical diversity library (**<u>Diversity Set</u>**) that identified a natural product chemotype that was next tested extensively for structure-activity relationships using single cell profiles of fundamental cellular processes (**Figure 1**). Here, the term ‘chemotype’ will be used to describe a set of molecules that share a molecular scaffold ^29–31^. The chemical family whose members most often displayed exceptional anti-leukemia bioactivity in initial tests with the **<u>Diversity Set</u>** was the rocaglates (flavaglines), a family of plant-derived translation inhibitors whose founding member, rocaglamide A (RocA), was identified for its anticancer activity ^32^. The Boston University Center for Molecular Discovery (BU-CMD) houses a series of synthetic rocaglate structural variants ^33^ which can be informally categorized into three subclasses: 1) regular rocaglates (RR), exemplified by natural products RocA (**<u>Supplementary Figure 1A</u>**), silvestrol, and eliptifoline, containing a cyclopenta[*b*]-benzofuran scaffold and no other ring fusions, 2) rocaglate pyrimidinones (RPs), exemplified by natural products aglaroxin C (**<u>Supplementary Figure 1B</u>**) and aglaiastatin, containing a mono- or bicyclic pyrimidinone ring system fused to the cyclopenta[*b*]-benzofuran core, ^34,35^ and 3) amidino-rocaglates (ADRs), a novel rocaglate subclass not found in nature, which contains an amidine ring fusion to the cyclopenta[*b*]benzofuran core and has been shown to exhibit the most potent translation inhibition for any rocaglate to-date ^36,37^. Following selection of rocaglates as a class for in-depth study, a representative **<u>Rocaglate Set</u>** of members, both natural and synthetic, of these rocaglate subclasses was assembled and deeply characterized for structure-activity relationships using a high dimensional single cell profiling panel.

**Figure 1.**
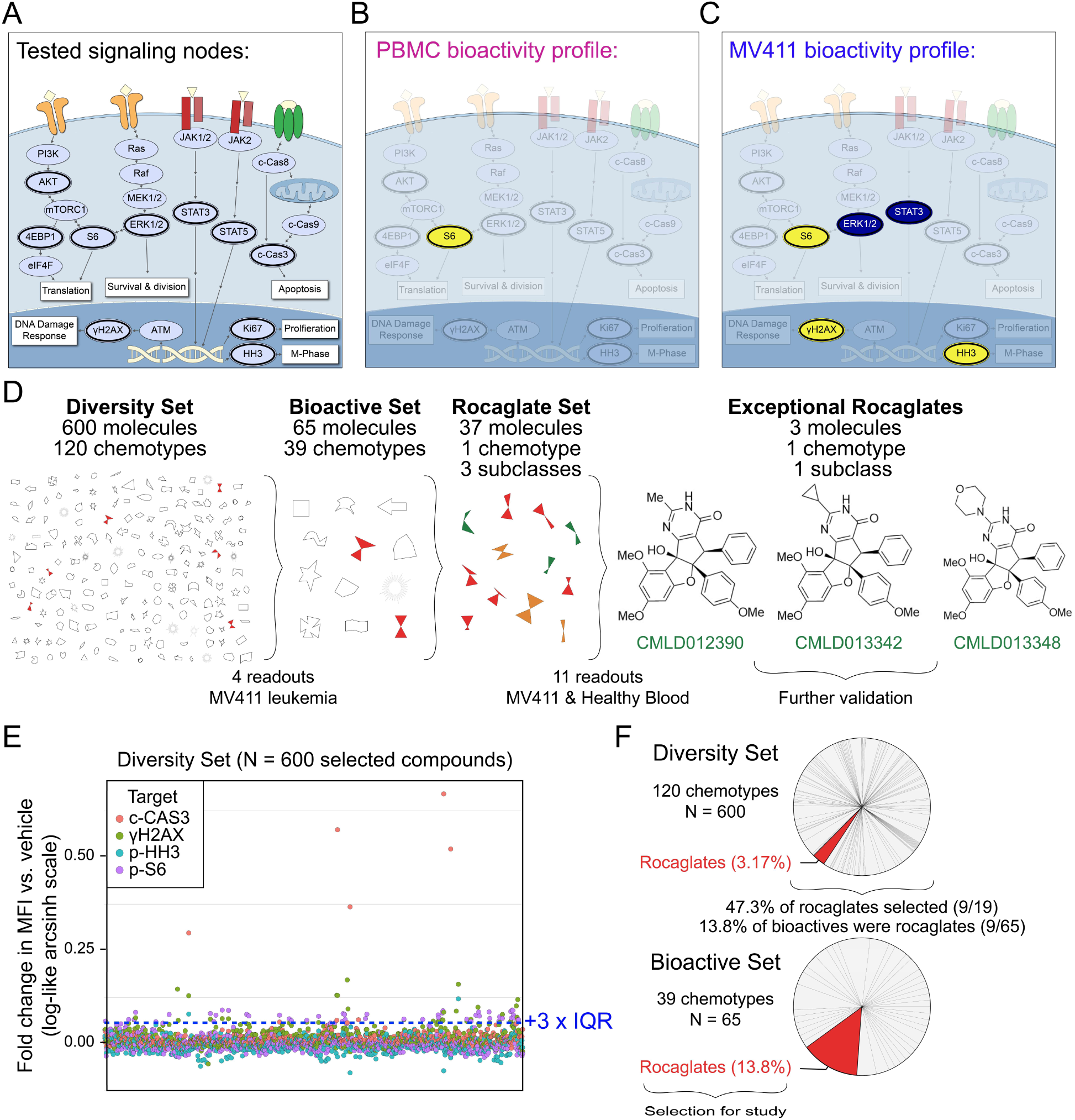
Rocaglates displayed exceptional activity in a multiplexed single cell bioactivity assay. **A)** A model of a cell annotated with the key readouts measured (dark outlines). mTOR pathway activation is explored through testing p-AKT, p-ERK1/2, p-S6, and p-4EBP1, which regulate translation, among other cellular processes. Activation of the DNA damage response is explored through γH2AX. Ki67 and p-HH3 are indicative of proliferation and M-phase of the cell cycle, respectively. p-STAT3 and p-STAT5 are transcription factors that regulate the expression of cell cycle, survival, and pro-inflammatory genes. Activation of apoptosis is measured through the detection of c-CAS3. Key readouts activated (yellow) or inhibited (blue) by the exceptional three rocaglates (shown in D) are depicted for PBMC in **B)** and MV411 in **C)**. **D)** Depiction of the progression from 600 compound **<u>Diversity Set</u>** to three exceptional rocaglates. A symbolic representation of the 600-compound **<u>Diversity Set</u>** comprised of 120 chemotypes, the 65 molecules identified as bioactive, and the **<u>Rocaglate Set</u>** selected for structure-bioactivity relationship studies. The names and structures for the three exceptional rocaglates are shown. The number of total molecules, chemotypes, and subclasses, where relevant, are depicted above each respective phase of testing. The number of readouts and cells used for testing are depicted below each phase. **E)** The arcsinh scaled fold change in median fluorescence intensity vs. vehicle for the 600-compound **<u>Diversity Set</u>** of natural products on the four readouts tested. Each readout is represented by a different colored circle. The bioactivity threshold was drawn at the vehicle median + 3 x interquartile range (IQR) (0.0573). **F)** The percentage of the 600 compounds from each of the 120 chemotypes represented in the **<u>Diversity Set</u>**. The pie slice corresponding to rocaglates is highlighted in red. Bioactive molecules were selected based on the threshold shown in **E)**. The percentage of the 65 bioactive molecules from each of the 39 chemotypes represented is shown, with the pie slice corresponding to rocaglates highlighted in red.

Ultimately, this work reveals subclasses and individual molecules with desirable and reproducible bioactivity in leukemia cells and not in healthy human blood cells. Additionally, this study shows the value of using single cell profiling to characterize a large network of fundamental cell signaling processes, as striking differences in bioactivity and cell type selectivity were observed between different rocaglate classes. In particular, rocaglate pyrimidinones (RPs) stand out as a subgroup that exclusively targeted leukemia cells and not healthy cells, with individual members displaying exceptional ability to activate intrinsic DNA damage responses that lead to the selective destruction of malignant cells. These exceptional RPs should now be prioritized in SAR and pre-clinical translational studies.

## Results

### Rocaglates are highly bioactive in leukemia cells

An initial test of 600 diverse molecules from the BU-CMD compound collection, with representation from 120 distinct chemotypes (**<u>Diversity Set</u>**), was conducted at 10 μM in MV411 leukemia cells using phospho-flow cytometry in combination with FCB for multiplexing. The goal of this experiment was to identify chemotypes with exceptional leukemia cell bioactivity. Exceptional activity following treatment with a compound or signaling input can be measured by phospho-flow in different ways, including exceptional potency (i.e., activity at nM or lower concentration)^38^, bioactivity (i.e., orders of magnitude greater responses in a measured proteins; e.g., high per-cell γH2AX indicating exceptionally strong DNA damage response)^16,18,38,39^, pathway selectivity or signature profile (i.e., a specific combination of activities per cell, such as activation of γH2AX without inhibition of p-4EBP1, indicating a DNA damage response without a halt to protein synthesis)^38,39^, and/or exceptional selectivity (i.e., targeting a specific subpopulation of cells, such as targeting cancer cells without targeting healthy cells)^16,38,40–42^. Here, the order of evaluation was first on bioactivity and selectivity, then on profile, and finally on evaluation of potency.

The panel selected for this initial test was designed to measure cell death, division, and translation and included post-translational modifications of proteins representing four fundamental cell processes: cleaved Caspase3 (c-CAS3, representing apoptosis), the gamma phosphorylation of H2AX (γH2AX, representing a DNA damage response), phosphorylated Histone H3 (p-HH3, representing M phase and cell cycle activity), and S235/S236 phosphorylated S6 (p-S6, representing active protein synthesis and cell growth) (**<u>Supplementary Table 1</u>**). The arcsinh fold change in median fluorescence intensity *vs*. vehicle was calculated for each compound across each of the four functional readouts (**Figure 1E**). For more details on standard arcsinh scaling using the inverse hyperbolic sine in cytometry analysis, see original use in the Supplement of ^41^ and recent implementations ^16,27,43,44^. While few compounds decreased the fold change in MFI *vs*. vehicle for any readout, 65 molecules were found to increase p-S6, c-CAS3, and γH2AX significantly compared with vehicle. Of the 65 bioactive molecules selected, 13.8% (9/65) were rocaglates, a chemotype which comprised only 3.2% of the initial set of 600, and predominant in the bioactive set (**Figure 1F**). Nearly half (9/19) of the rocaglates tested in the **<u>Diversity Set</u>** were selected as bioactive. Of the bioactive rocaglates, driving activities included c-CAS3 and γH2AX. Based on their high frequency of triggering c-CAS3 and/or γH2AX in MV411 leukemia cells, rocaglates were prioritized for further investigation.

All 19 rocaglates in the initial 600 compound **<u>Diversity Set</u>** were RRs. To investigate whether rocaglates from different structural subgroups were bioactive in MV411 leukemia cells and whether this bioactivity extended to healthy cells, we tested an expanded library of 37 rocaglates, intentionally weighted toward known bioactive molecules, with representation from each of the three subclasses at 10 µM on MV411 and healthy peripheral blood mononuclear cells (PBMCs) (**Figure 2A**). The primary staining panel was selected to test a range of cell functions including cell growth, translation, and death and has been validated in previous work: Ki67 (cell proliferation ^45,46^), phospho-S6 ribosomal protein at serine 240/244 (p-S6 S240/244) (growth, AKT/mTORC1 specific ^47,48^), phosphorylation of gamma H2AX at serine 139 (γH2AX) (DNA damage response ^49^), phosphorylation of STAT3 at serine 727 (p-STAT3) (amplified transcriptional activity, downstream mTOR ^50^), phosphorylation of STAT5 at tyrosine 694 (p-STAT5) (cell survival, highly phosphorylated in cancer cells ^51,52^), phospho-S6 ribosomal protein at serine 235/236 (p-S6 235/236) (growth, activated by ERK/RSK and AKT/mTORC1 ^47,48^), phosphorylation of ERK1/ERK2 at threonine 202, tyrosine 204 (p-ERK1/2) (proliferation, activated by MAP kinase cascade ^53,54^), phosphorylated histone H3 at serine 28 (p-HH3) (growth, cell cycle M phase ^55^), phosphorylation of 4EBP1 at threonine 37/46 (p-4EBP1) (growth, activated by mTORC1 ^56^), phosphorylation of lymphocyte-specific protein tyrosine kinase at tyrosine 505 (p-LCK) (SRC-family kinase involved in T-cell receptor signaling ^57^), and phosphorylation of Akt at serine 473 (p-Akt) (Upstream of mTOR ^41,58^) (**<u>Supplementary Table 1)</u>** _59,44,60._

**Figure 2.**
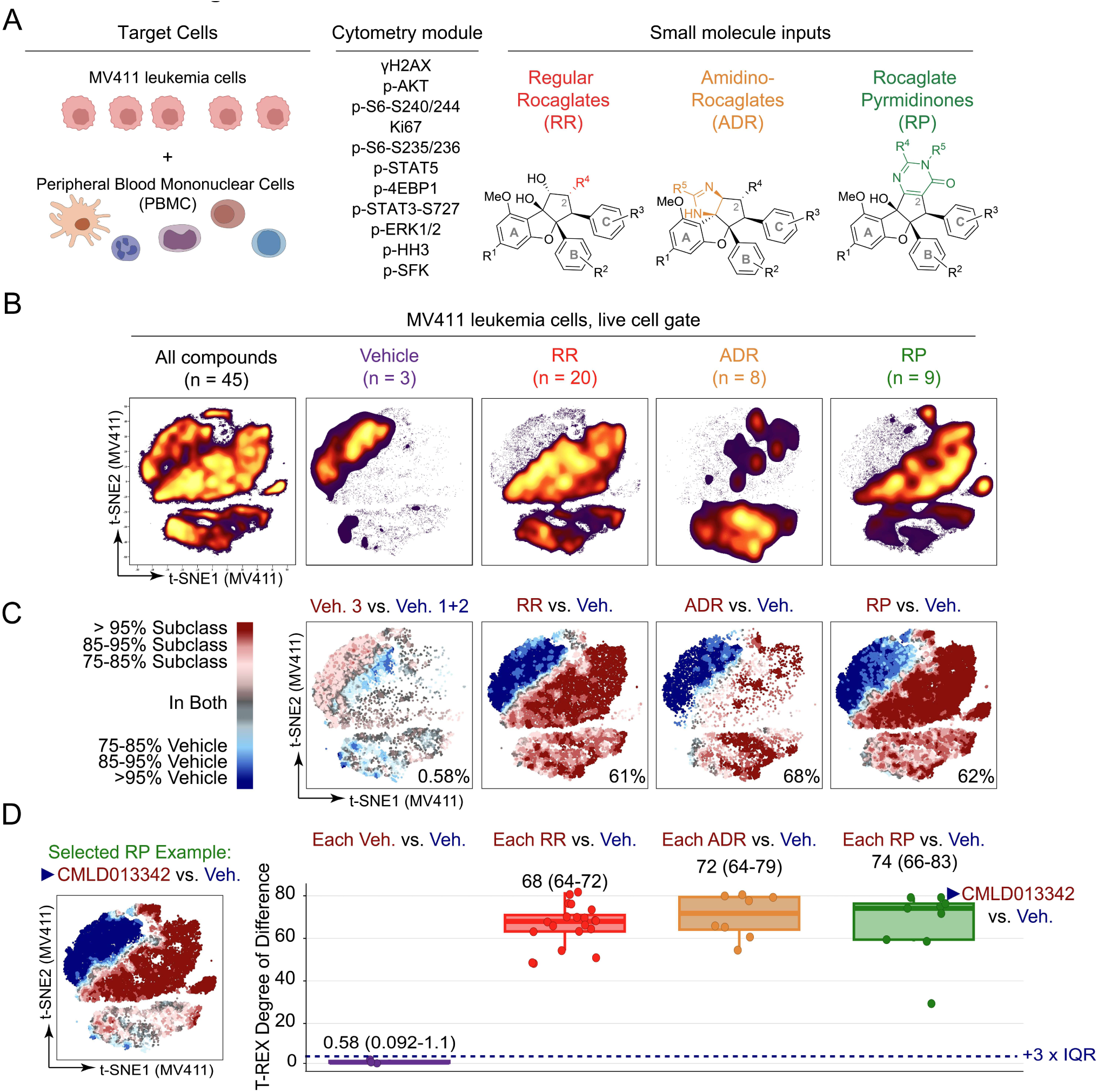
Rocaglates are a chemical family with bioactivity against leukemia cells. **A)** Shown, are the three elements core to a single cell chemical biology experiment conducted using flow cytometry – target cells, a module of cytometry readouts, and small molecule inputs - and the inputs selected for these studies. The MV411 leukemia cell line and human peripheral blood mononuclear cells were selected as the set of target cells. Eleven readouts of core cell functions were selected for measuring per cell as part of the cytometry module. Lastly, rocaglates falling into the three depicted structural subclasses – regular rocaglates, amidino rocaglates, and rocaglate pyrimidinones – were chosen as the small molecule inputs. **B)** Plot depicting the result of performing a t-SNE analysis on the entire pre-processed MV411 dataset (All compounds). The t-SNE of all compounds is divided based on the category of input (Vehicle, RR, ADR, and RP) and colored based on cell density. **C)** T-REX plots depict regions of significant difference between the t-SNE of one rocaglate subclass vs. vehicle-treated cells in MV411. As a representative of variation between the three tested vehicle wells, a T-REX plot is shown for the analysis of vehicle 3 vs. the pooled set of vehicles 1 and 2 (leftmost plot). The T-REX degree of difference [(number of red cells + number of blue cells)/total number of cells] for each analysis is reported in the lower right corner of each plot. **D)** Box and whisker plot of T-REX degree of difference for each compound vs. vehicle-treated cells in MV411. Corresponding box and whiskers are grouped and colored according to rocaglate structural subclass. The median and IQR range of the T-REX degree of difference for each subclass is provided above the respective set of box and whiskers. The dashed navy line is shown at the vehicle median + 3 x IQR to indicate a significance threshold. Representative T-REX plot for **CMLD013342** vs. vehicle is shown to the left of the box and whisker plot for reference on the analysis conducted. T-REX plots for the remaining compounds are depicted in **<u>Supplementary Figure 2B</u>.**

Next, the t-SNE algorithm was used to reduce the dimensionality of the data from 11-dimensional space to 2-dimensional space for visualization and analysis. The t-SNE plot was made using MV411 data from all 45 conditions tested (37 rocaglates, 3 vehicles, and 5 controls which are not shown) and then subdivided based on the subclass of origin (and vehicle) for comparison of high dimensional signaling profiles (**Figure 2B)**. Striking differences were apparent in t-SNE plots for each of the three rocaglate subclasses when compared to the pooled vehicle data: vehicle cell density was localized to the upper left side of the map, whereas cell density for the rocaglate subclasses was nearly absent from this region. These differences were quantified by T-REX. Specifically, the t-SNE from each of the three respective rocaglate subclasses was compared to the pooled set of the 3 vehicle wells (**Figure 2C**). The T-REX plots revealed the cell populations enriched in vehicle-treated cells (blue) and rocaglate subclass-treated cells (red). For each T-REX plot, the T-REX degree of difference [(number of red cells + number of blue cells)/total number of cells] was computed to quantify how different subclass-treated cells were from vehicle-treated cells. All rocaglate subclass-treated cells had a T-REX degree of difference that was greater than 60%, indicating substantial change upon 16 hours of rocaglate treatment, relative to vehicle. In comparison, when T-REX was performed on vehicle 3 *vs*. the pooled set of vehicles 1 and 2, the T-REX degree of difference was <1%, indicating the vehicles were highly similar to each other. The contrast in cell density apparent in t-SNE and quantified by T-REX arises from phenotypic differences in MV411 cells treated with molecules from the three subclasses, as compared to vehicle, and confirms the original observation (**Figure 1**) that the three rocaglate subclasses display outstanding activity against leukemia cells.

To investigate the variation in bioactivity within each subclass, T-REX was applied by comparing the t-SNE from each compound to the pooled set of the 3 vehicle wells. As a representative of this analysis, the T-REX plot for **CMLD013342** vs. vehicle is shown in **Figure 2D** (data for remaining compounds are shown in **<u>Supplementary Figure 2</u>**). Then, a T-REX degree of difference calculation was performed for each of the 40 comparisons (37 rocaglates + 3 vehicles). All 37 rocaglates had a T-REX degree of difference above a threshold of three times the interquartile range of the vehicle data over the vehicle median (+3 x IQR). Taken together, these results indicated that all tested rocaglates were bioactive against MV411 leukemia cells.

### Rocaglates have distinct, subclass-specific bioactivity patterns in leukemia and healthy blood

To further explore signaling patterns within and across the rocaglate subclasses, the field standard log-like arcsinh ratio of the median phospho-protein signal in treated cells as compared to the signal in the first vehicle *(i.e.*, arcsinh [MFI_treated_/MFI_vehicle1_]) was performed for each marker and compound. This resulted in values that range from 0 (no change) to +1 arcsinh unit (an increase of X-fold, yellow) or -1.5 (a decrease of 1.5X-fold, blue). In **Figure 3A**, compounds were arranged according to rocaglate subclass to allow comparison of phenotypic differences within and between subclasses. In **Figure 3B**, the same data were arranged by hierarchical clustering of the signaling profile (below). When organized by subclass, RPs displayed a more consistent signature than other subclasses (median IQR across all markers for RP = 0.07, RR = 0.2, and ADR = 0.2). The RP signature, which was also observed in some RRs, included exceptionally strong γH2AX and inhibition of p-ERK, a signature indicating activation of DNA damage response and inhibition of MAPK proliferative signaling. In contrast to RPs, most ADRs inhibited γH2AX and p-4EBP1 and activated p-AKT and p-STAT5, a signature suggesting activation of cell survival signaling that would be undesirable in an anti-cancer compound. RRs were heterogeneous in the pattern of phosphorylation seen following rocaglate treatment and had members that displayed patterns that were similar to either RPs or ADRs. Grouping by unsupervised clustering of bioactivity led to the successful separation of RPs into clusters 2 and 3 and ADRs into clusters 4 and 5 (except for **CMLD012600**); RRs were interspersed throughout clusters 2 - 5. One readout that strongly contrasted between clusters was γH2AX; while cluster 3 had the highest γH2AX compared to vehicle, cluster 2 had moderately high γH2AX compared to vehicle, cluster 5 had similar γH2AX compared to vehicle, and cluster 4 had low γH2AX compared to vehicle. Structures for the 37 rocaglates grouped according to rocaglate subclass and hierarchical clustering can be found in **<u>Supplementary Figure 3</u>** and **<u>Supplementary Figure 4</u>**, respectively.

**Figure 3.**
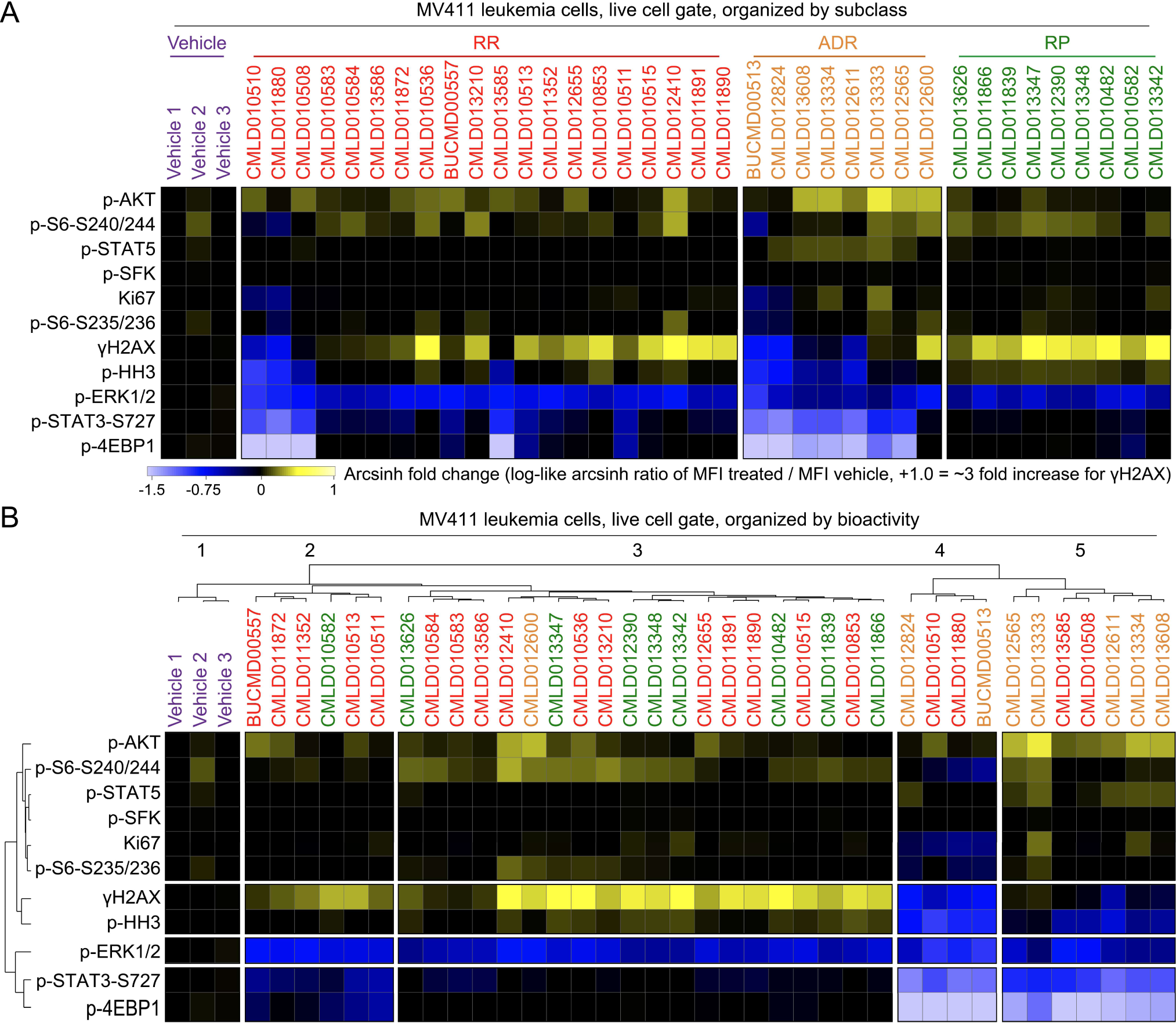
Rocaglate subclasses had distinct patterns of bioactivity. **A)** Heatmap depicting the arcsinh ratio of the median fluorescence intensity for each compound (listed on top of heatmap) and readout (listed left of heatmap) by the median fluorescence intensity of Vehicle 1. Cells on the heatmap range from light blue for the lowest values to bright yellow for the highest values. Compounds are grouped and colored according to the rocaglate subclass listed on the top of the plot. **B)** Heatmap as in Figure 2B clustered according to a dendrogram of the transformed median fluorescence intensity for each compound and readout.

To investigate how γH2AX induction patterns by rocaglates compared between MV411 leukemia cells and healthy blood, the percentage of γH2AX positive (% γH2AX+) cells for each compound (and vehicle) was quantified for each cell type (**Figure 4A**). A threshold of 15% γH2AX+ cells was selected to distinguish compounds that trigger significant DNA damage in MV411 and PBMC. Only the vehicles and three RRs did not trigger significant DNA damage in either cell type. Nearly all ADRs and one RR (**SDS-1-021** ^61–65^, represented twice in the set as both racemic (**CMLD011880**) and enantioenriched (**CMLD010508**) stocks) triggered significant DNA damage in PBMC alone. Distinctly, two ADRs induced significant DNA damage in both cell types. Lastly, all RPs and some RRs led to leukemia-specific induction of γH2AX. Quantification of γH2AX induction in both cell types further illuminated rocaglate structural subclass bioactivity signatures and outlier compounds within each subclass.

**Figure 4.**
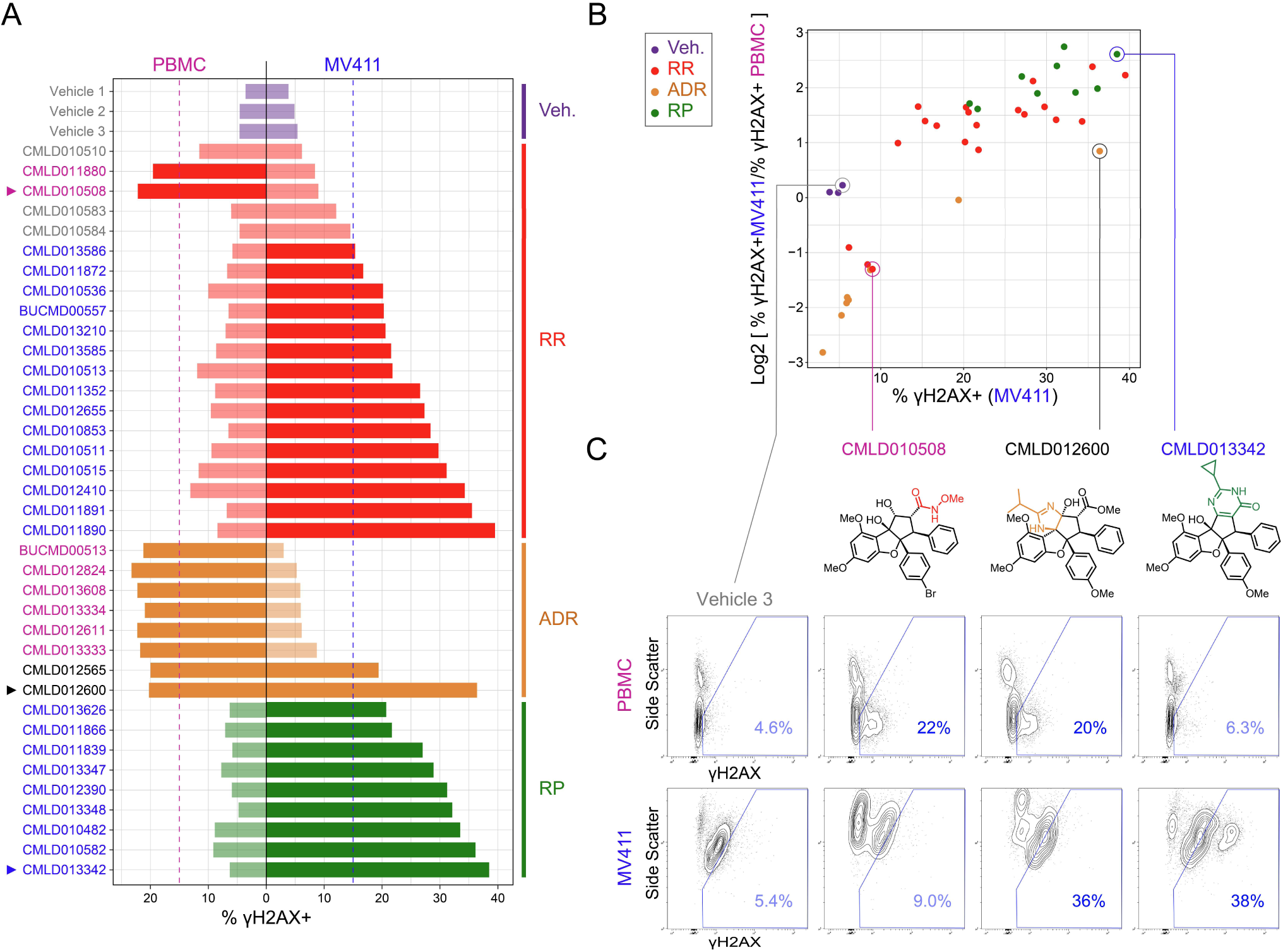
MV411 specific activity was observed in the RP subfamily and some regular rocaglates. **A)** Bar plot of percentage of γH2AX positive (% γH2AX +) cells for each compound grouped and colored according to rocaglate subclass listed on the right side of the plot. Bars on the left side of the vertical black line correspond to the compound response in peripheral blood mononuclear cells (PBMC) and the right side corresponds to the compound response in MV411 cells. The dotted vertical blue line corresponds to a significance threshold for % γH2AX+ cells in MV411 and the dotted pink line corresponds to a significance threshold for % γH2AX+ cells in PBMC. Compounds that cross the threshold have darkened colored bars. The compound name (or vehicle) is listed on the far left side of plot colored according to the following system: blue = increases % γH2AX+ cells past threshold in MV411 and not in PBMC, pink = increases % γH2AX+ cells past threshold in PBMC and not in MV411, light grey = does not increase % γH2AX+ cells past threshold in either cell type, dark grey = increases % γH2AX+ cells past threshold in both cell types. Arrows are displayed to the left of compounds that will be shown in Figure 4C. **B)** Scatter plot of % γH2AX+ cells in MV411 on the x-axis and the log2 fold ratio of % γH2AX+ cells in MV411 to % γH2AX+ cells in PBMC on the y-axis. Each dot corresponds to a compound colored according to rocaglate structural subclass. Compounds that will be shown in Figure 4C are circled and labeled. **C)** Contour plots of Side Scatter vs. γH2AX for Vehicle 3, CMLD010508, CMLD012600, and CMLD013342, respectively from left to right in PBMC (top) and MV411 (bottom). The blue line indicates the γH2AX+ gate. The % γH2AX+ cells within the gate is written in blue in the lower right corner. Corresponding structures for each compound are depicted below the compound name.

To identify compounds with desirable anti-leukemia activity, the compounds with the greatest leukemia-specific induction of DNA damage were next identified. The log2 fold ratio of the percentage of γH2AX+ cells was compared between MV411 and healthy PBMC cells (i.e., log2 [% γH2AX^+^_MV411_/ γH2AX^+^_PBMC_], **Figure 4B**). The ADRs that had PBMC-specific induction of DNA damage in **Figure 4A** were in the lower left quadrant, as they had the lowest leukemia-specificity of DNA damage induction and low induction of DNA damage in MV411. All RPs and the select RRs that were demonstrated to have leukemia-specific induction of γH2AX in **Figure 4A** were found to be in the top right quadrant, as they had maximal DNA damage induction in MV411 and a specificity of this induction to leukemia cells. Three RPs, **CMLD012390**, **CMLD013342**, and **CMLD013348** had the highest log2fold ratio of % γH2AX+ in MV411 to % γH2AX+ PBMC. The representative contour plot of **CMLD013342** depicted in **Figure 4C** demonstrated a distinct population of γH2AX hyperactivated cells that was not seen in the contour plot for **CMLD012600**, a rocaglate with a similar % γH2AX+ cells. This γH2AX hyperactivated population of cells by **CMLD013342** was also not present in the PBMC contour plot. The uniform induction of a leukemia-specific DNA damage response associated with cell death and γH2AX hyperactivated population of cells, helped establish RPs as an interesting subclass of rocaglates for further investigation as potential leukemia therapeutics.

### The rocaglate RP subclass selectively induces a γH2AX+ p-4EBP1+ bioactivity pattern seen only in leukemia cells

To visualize the differences in high-dimensional signaling profiles across rocaglate subclasses, a T-REX analysis was performed by comparing the t-SNE from each of the three respective rocaglate subclasses to the pooled set of the other rocaglate structural subclasses (including vehicle) (**Figure 5A**). These T-REX plots identified the cell populations that were enriched in (red) and absent from (blue) a given subclass. Notably, T-REX illuminated a subpopulation of cells specifically induced by RPs (RP island). Coloring the grouped t-SNE for all 45 compounds based on measurements for each functional readout suggested that the RP island was a region of cells with high median fluorescence intensity of p-4EBP1 and γH2AX (**<u>Supplementary Figure 5</u>**).

**Figure 5.**
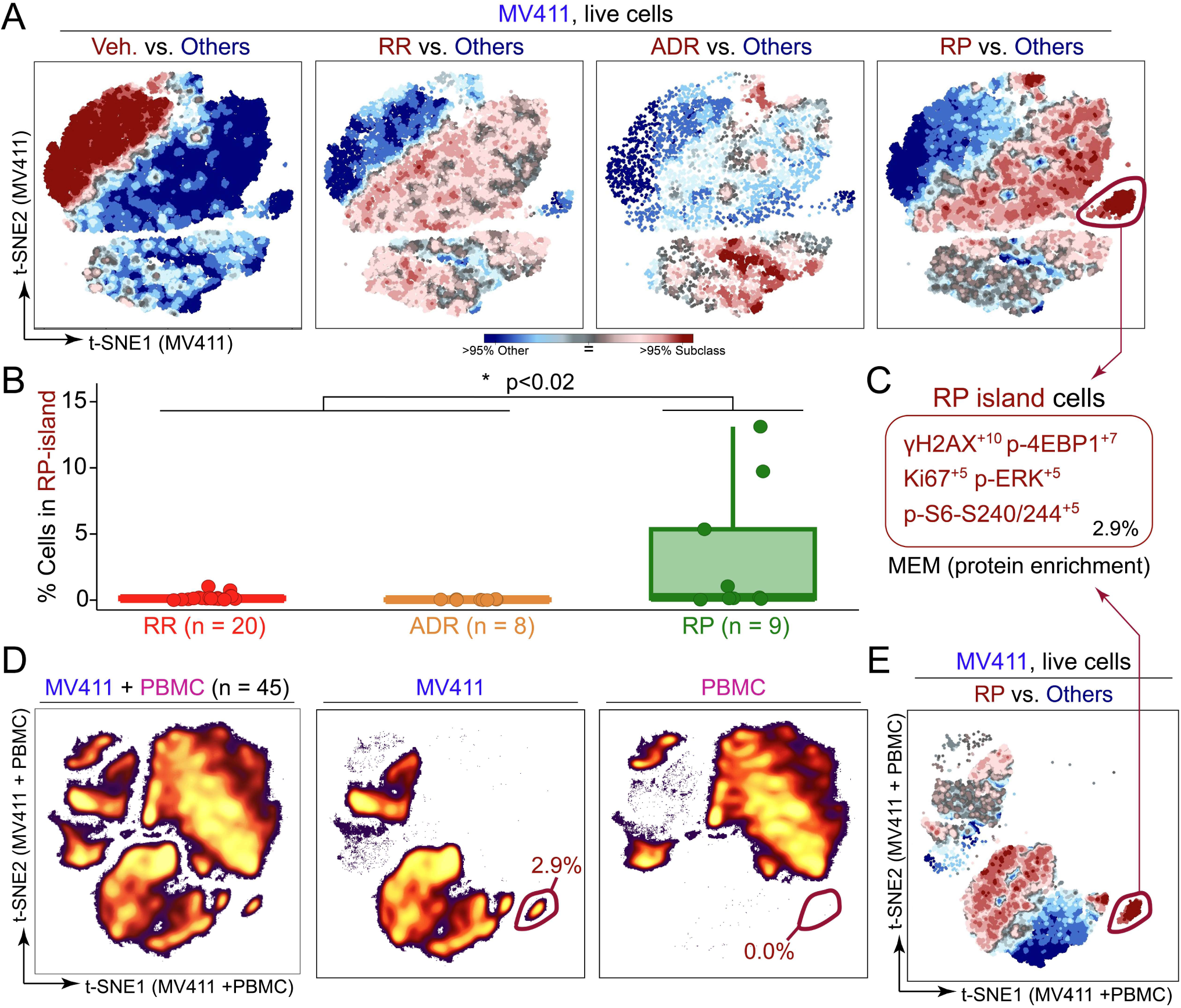
The RP subfamily had a distinct, leukemia-specific population of cells with high DNA damage response. **A)** T-REX plots depict regions of significant difference between the t-SNE of one rocaglate subclass vs. the pooled set of all three remaining subclasses (including vehicle) for MV411 data. The RP island is circled in red in the “RP vs. Others” plot. **B)** Boxplot depicting the percent of cells in the RP island for MV411 for each rocaglate grouped by rocaglate subclass. The Wilcoxon rank sum test of RP vs. remaining subclass % in gate was conducted and generated a p-value of 0.015. **C)** MEM protein enrichment label generated for the RP island of cells. The percentage of the total population of cells for the RP island is depicted in the lower right corner. **D)** t-SNE plot of the MV411 and PBMC data on a new embedding with corresponding x and y axes (left). The t-SNE plot for both cell types was divided into separate plots for MV411 (middle) and PBMC (right). The percentage of cells in an MV411-specific island is circled in red and depicted in the lower right of the MV411 and PBMC plots. **E)** T-REX plot of RPs vs. the pooled set of all three remaining subclasses (including vehicle) for the MV411 data using the t-SNE performed on the MV411 and PBMC data together shown in Figure 5D. The MV411-specific island circled in red was identified to have the same MEM label as the RP island.

To determine the true specificity of this island to RPs, the percentage of cells in the RP island for each rocaglate was calculated (**Figure 5B**). A Wilcoxon rank sum test of the percentage of cells in the RP island for RPs vs. the remaining subclasses was conducted as a test of significance and generated a p-value of 0.015. The percentage of cells falling into the RP island was also correlated with transformed 98^th^ percentile fluorescence intensity for γH2AX and p-4EBP1 (**<u>Supplementary Figure 6</u>**). A Marker Enrichment Modeling (MEM) ^28^ analysis was performed on the RP island to characterize contextual protein enrichment (absolute MEM labels range from 0 = no enrichment to +10 = highest enrichment). The resulting MEM label was: γH2AX+10, p-4EBP1+7, Ki67+5, p-ERK+5, and p-S6 - S240/244 +5 (**Figure 5C**). The enrichment for γH2AX and p-4EBP1 indicated that RP island cells had exceptionally high bioactivity, seen in the activation of the DNA damage response (γH2AX), and a distinct signature, seen in retention of some mTOR pathway activity (p-4EBP1).

To assess the leukemia-specificity of the RP-island, as activation of the DNA damage response in healthy cells would be detrimental, a t-SNE analysis was performed on the MV411 and PBMC together to generate a new embedding with corresponding x and y axis scales (**Figure 5D**). After dimensionality reduction with t-SNE, cells were separated based on origin (MV411 or PBMC) (**Figure 5D**). The MV411 t-SNE analysis revealed a distinct island of cells from the remaining cell density on the lower right side of the plot. When quantified, this cell population was >99% specific to MV411. This leukemia-specific island was confirmed as the RP island after a MEM analysis generated the same protein enrichment label as displayed in **Figure 5C** (**Figure 5E**). Together, these results established that RPs activate a strong and leukemia-specific DNA damage response through the RP island.

### RP structure correlates with exceptional DNA damage responses in leukemia cells

While the Wilcoxon rank sum test provided a subclass-level validation of the RP specificity of the RP island, further investigation of bioactivity and structure at the compound level was necessary for understanding which compounds shifted cells to the RP island and why. To address this, the t-SNE map of pooled RPs was further subdivided based on compound of origin, and the percentage of total cells falling into the RP island was quantified for each RP (**Figure 6).** This compound-level analysis highlighted three RPs, **CMLD012390**, **CMLD013342**, and **CMLD013348**, as the only compounds with greater than 5% of cells in the RP island gate. Markedly, these are the same compounds with the highest log2fold ratio of % γH2AX+MV411/% γH2AX+ PBMC as reported in **Figure 4B**. These three compounds commonly possess three unique structural features that are found in isolation in the other six RPs, specifically a non-aryl pyrimidinone ring substituent (termed the RP “R-group”), a 4′-methoxy substituent on the rocaglate “B”-ring, and a monocyclic, *N*-unsubstituted pyrimidinone ring. This suggested that a combination of multiple structural features might drive SAR for the RPs with high cell density in the RP island. Thus, the RP island was associated with an exceptional, leukemia-specific induction of the DNA damage response and prominent structural features were identified for future SAR studies.

**Figure 6.**
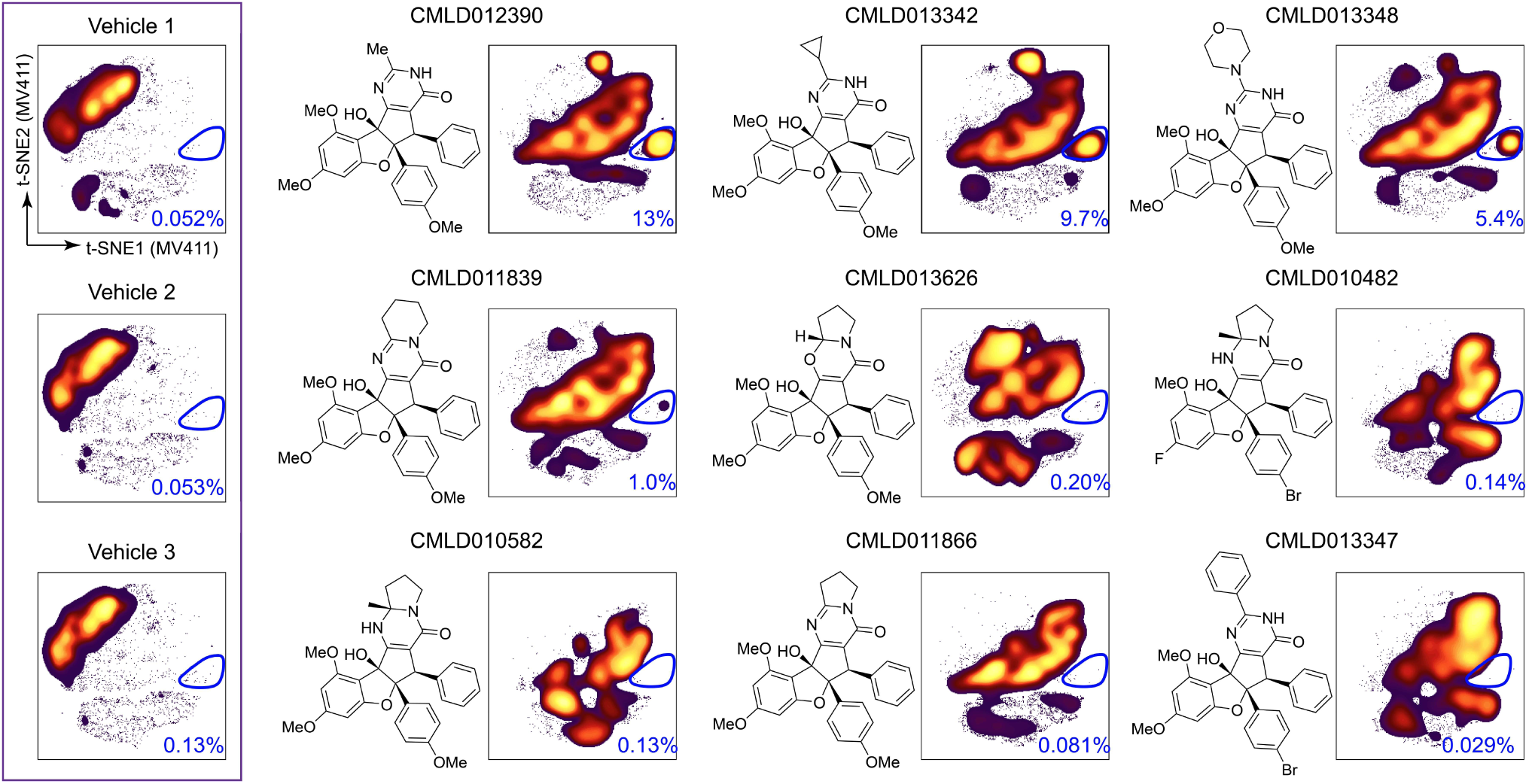
Rocaglates contributing to RP-island possessed structural commonalities. t-SNE shown in Figure 2B of all compounds in MV411 divided based on rocaglate well of origin is shown for the set of 9 RPs in order of decreasing percentage of cells in the RP island (t-SNEs for all individual compounds are shown in **<u>Supplementary Figure 2A</u>**). The t-SNE plots for the three vehicle wells are shown for reference in the purple box on the left. The percentage of cells in the RP island gate circled in dark blue is included at the bottom right of each plot. The chemical structure corresponding to each rocaglate is depicted on the left side of each t-SNE plot (except the three vehicle wells).

To further investigate the potency and timing of γH2AX activation by these special RPs, dose-response and time course experiments were performed in MV411 cells. For the dose-response, six doses (10, 5, 1 0.1, 0.05, 0.01, and 0 µM) were selected to encompass a 1000-fold range and the top two RPs, **CMLD012390** and **CMLD013342** were chosen for testing. A γH2AX + gate was drawn on biaxial plots of Side Scatter Area (SSC-A) vs. γH2AX based on γH2AX levels in the vehicle-treated cells. The percentage of cells within the γH2AX + gate was quantified for each concentration and used to calculate the half-maximal activating concentration (EC_50_) **(Figure 7A**). **CMLD012390**, the RP with the greatest percentage of cells in the RP island in **Figure 6**, had an EC_50_ of 213 nM, and **CMLD013342**, the RP with the second greatest percentage of cells in the RP island had a slightly higher EC_50_ of 492 nM. **CMLD012390** and **CMLD013342** were also used to challenge MV411 cells at 1, 4, and 16 hours to investigate the timing of γH2AX induction. Within 4 hours, mild γH2AX induction was seen for both compounds; this γH2AX induction increased in potency to match levels seen in both the **<u>Rocaglate Set</u>** and dose-response within 16 hours (**Figure 7B** and **<u>Supplementary Figure 7</u>**). Thus, these two RPs shared multiple structural commonalities and demonstrated a strong and reproducible induction of DNA damage response within 4 hours. Going forward, future SAR studies should involve direct, pairwise comparisons to discern the individual impact of the unique structural features in these RPs on specificity in selectively inducing DNA damage responses in leukemia cells and not in healthy blood.

**Figure 7.**
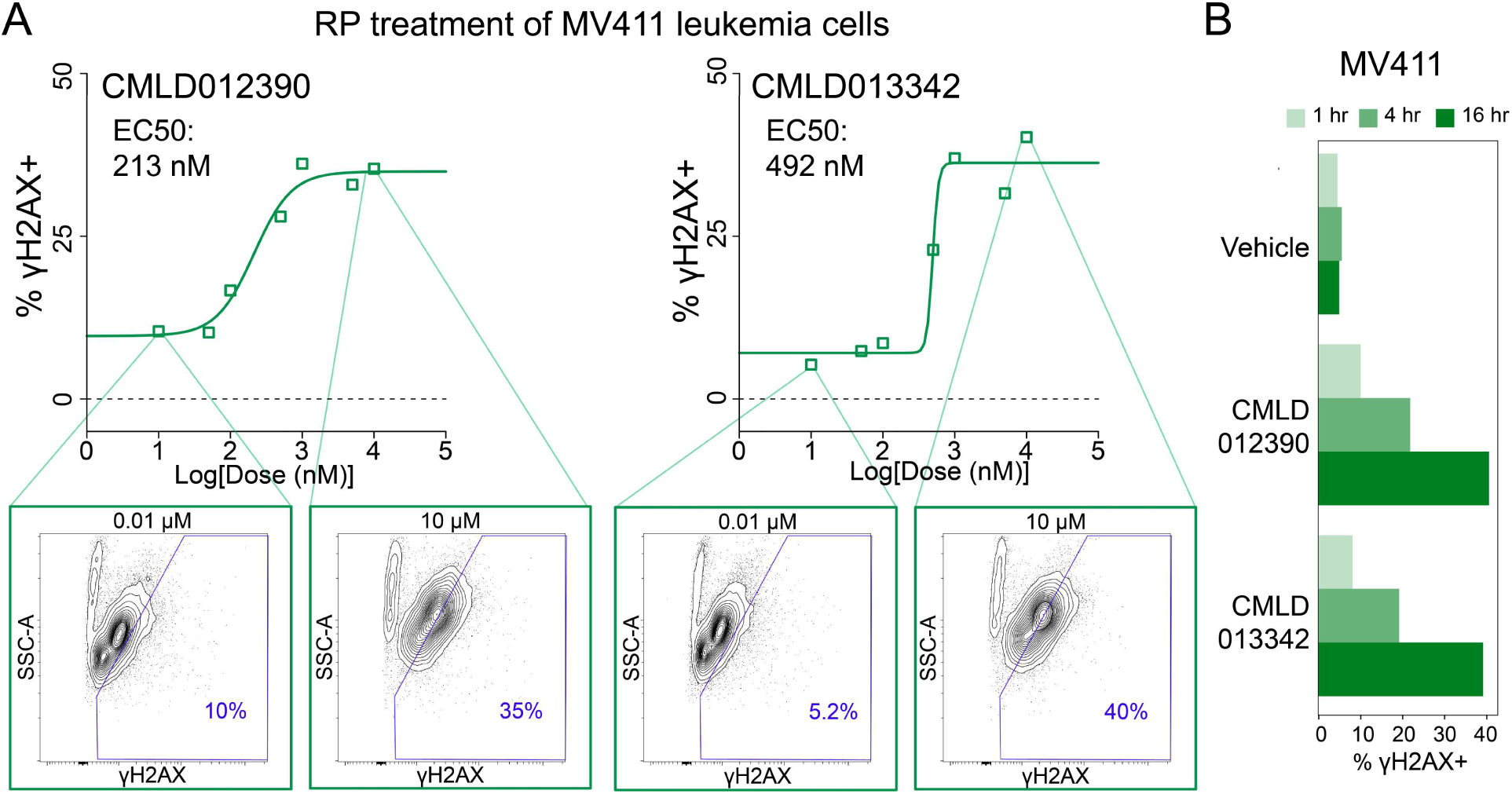
The top 2 RPs activated γH2AX with < 500nM potency within 4 hours. **A)** Dose-response titration curves depicting % γH2AX+ cells (based on expert gating) vs. log[dose(nM)]. A half-maximal activating concentration (EC50) curve was fitted to the data and γH2AX EC50 values are shown in the top left of each plot. Contour plots of Side Scatter Area (SSC-A) vs. γH2AX at 10 µM and 0.01 µM, respectively, with 10% of cells per contour are shown for each compound. A polygon gate is drawn in blue to delineate γH2AX+ cells, with the % γH2AX+ cells within the gate depicted in the lower right corner. **B)** Bar plots depicting % γH2AX+ cells derived from a time course experiment at 1, 4, and 16hr for Vehicle, **CMLD012390**, and **CMLD013342**.

## Discussion

An advantage of single cell chemical biology approaches is the potential to test a greater space of cell biology readouts and to do so with primary cells that include diverse healthy cell types. Thus, phospho-flow and single cell chemical biology approaches can be used to include a range of cells that will be present in the human tissues in which compounds will need to function *in vivo*. Here we report a test of diverse compounds facilitating structure-activity relationship studies based on multiparameter, single cell phospho-flow assays. This revealed rocaglates as a highly bioactive chemotype and identified all rocaglate pyrimidinones tested and most amidino-rocaglates tested as chemotypes that respectively selected for leukemia or healthy blood cells. The comparison between primary human blood and leukemia cells acts as a proof of concept for potentially employing human tissues in early phase SAR studies.

Since phospho-flow was first combined with FCB for chemical biology screening ^6^, this approach has been used for screening and characterization of promising compounds based on cell type- and pathway-specific effects. The screen of small molecules described here was similarly performed in combination with FCB for multiplexing and used a focused panel of four functional states of proteins to efficiently profile the modulation of relevant pathways involved in leukemia cell translation and death. The screening set of 600 structurally diverse compounds is a relatively small library compared to what might be possible if this approach were scaled; part of the success of the study with this focused library is likely due to the selection of key functional signaling events to measure as bioactivity readouts. Here, multiplexed cytometry measured 2,400 cell-based biological readouts at the single cell level in one experiment. Future studies might go beyond this work by leveraging additional advantages of phospho-flow signaling profiles, especially the ability to resolve impacts on different subpopulations of primary cells in blood, tumors, or other tissue. Sixty-five promising activators were selected for further investigation based on activity at p- S6, here interpreted as measuring growth and translation, c-CAS3, an indicator of apoptosis, and/or γH2AX, a readout of DNA damage. While previous studies have focused on selection of individual hits for follow-up testing, this approach was focused instead on identifying promising chemotypes. Of the 65- compound bioactive set, 9 of the compounds were rocaglates, a natural product family that has received a great deal of attention for promising anti-cancer and anti-infective activities.

While rocaglate bioactivity has been of interest since rocaglamide A (RocA), the prototypic rocaglate, was first isolated in 1982, this study was the first phospho-flow-based test of rocaglates and one of only a few that have characterized the bioactivities of various structural subclasses within the chemotype ^32^. Larger screens of >200 structurally diverse rocaglates have mainly monitored clamping of the three eIF4A homologs (eIF4A1-3) to RNA, *in vitro* translation, and cytotoxicity ^36,37,66^. This test set of 37 rocaglates from three structural subclasses profiled bioactivity using 11 functional states of proteins across MV411 and PBMC cells, enabling 22 cell-based biological readouts to be tested for each compound simultaneously at the single cell level.

Cytometric profiling of the 37 rocaglates in combination with t-SNE and T-REX enabled us to establish that these molecules were all bioactive against leukemia cells, considering all 11 protein functional readouts measured at the single cell level. Additionally, investigation of fluorescence intensity for each molecule and readout enabled rocaglate bioactivity within and across subclass to be compared. There were noticeable similarities in fluorescence intensity for readouts within each subclass and differences across subclass: RPs were distinguished by high γH2AX, p-S6 S240/244, and p-HH3 and ADRs demonstrated low γH2AX and p-4EBP1 and high p-AKT and p-STAT5. However, a dendrogram of transformed fluorescence intensity across all 11 functional readouts and 37 compounds only separated ADRs from RPs; this suggested that the previously identified structural subclasses may not fully capture bioactivity differences.

Analyzing γH2AX, a readout that contrasted across the clusters of rocaglates formed in **Figure 3**, in both PBMC and MV411 shed even further light on bioactivity differences between structural subclasses. While some RRs and all RPs activated γH2AX in MV411 and not PBMC, ADRs only activated γH2AX in PBMCs. For RRs these findings are not surprising; rocaglamide A, an RR, has demonstrated leukemia-specific bioactivity in previous studies ^67^. However, ADRs have been found to be the most potent *in vitro* RNA clampers of eIF4A1 and eIF4A2 ^35,37^. Interestingly, RPs, the subfamily demonstrating uniform leukemia-specific induction of γH2AX, have primarily been reported as agents for hepatitis C virus (HCV) and to show a possible bias for the inhibition of viral entry over translation inhibition ^34^. The natural product aglaroxin C (**CMLD011866**) has been cited as a potent cytotoxic agent against multiple cancer cell lines, though not yet as selective for cancer cells as opposed to healthy cells ^68–71^. Therefore, these findings reveal leukemia-specific bioactivity, or a lack thereof, for RPs and ADRs, respectively.

By leveraging t-SNE, T-REX, and MEM, we were able to identify a small subpopulation of cells with inflated γH2AX that was only present in RP-stimulated MV411 cells. Given that this population was <3% of total MV411 cells, it is likely that this would have been overlooked by approaches only accounting for bulk cell responses. Further investigation allowed us to uncover **CMLD012390**, **CMLD013342**, and **CMLD013348** as the primary contributors to this island of cells. This response was not seen by aglaroxin C (**CMLD011866**) though its potent cytotoxic activity is warranted based on prior literature ^69,70^. γH2AX is produced in response to histone H2AX phosphorylation on a serine four residues from the carboxyl terminus and acts as a sensitive marker for DNA double-stranded breaks (DSBs). Many existing cancer therapies such as etoposide and mitoxantrone act by introducing sufficient DSBs to activate cell death pathways.^72^ Therefore, the ability of these top three RPs to potently activate γH2AX in only leukemia cells should now be studied for potential clinical use. Rocaglates have been shown to have two key classes of DEAD-box RNA helicase targets: the three eIF4A homologs, and more recently DDX3 ^73,74^. While eIF4A is implicated in translation initiation, DDX3 is involved in a range of functions related to RNA metabolism^75^.

The compounds **CMLD012390**, **CMLD013342**, and **CMLD013348** have not been previously reported as distinct members of the RP subfamily. Notably, these compounds possess some structural similarities; they commonly have a methoxy group on their B-ring (the southeastern ring), a non-aryl R-group (the northern projection from the pyrimidinone), and a monocyclic, N-unsubstituted pyrimidinone with unique steric properties that, in contrast to *N*-substituted RPs such as **CMLD011866**, is capable of equilibrating to an aromatic hydroxypyrimidine tautomeric form. Further structure-activity relationship and target identification studies may be used to confirm which structural commonalities possessed by the top three RPs drive their exceptional γH2AX activation ability. While the RP-specific γH2AX signature (the RP island) for **CMLD012390**, **CMLD013342**, and **CMLD013348** may imply an as-yet unknown target, it is also possible that differences in potency or timing of engagement with eIF4A1 and/or DDX3 drive this unique activity signature.

In this study, we investigated structure-activity relationships for 37 rocaglates across 11 key functional proteins representing hallmarks of cell biology in MV411 leukemia cells and PBMCs. The functional states of proteins chosen, while more expansive than a single target- or readout-based approach, did not capture all potentially impacted pathways. For confirming the leukemia-specificity of the bioactive molecules identified, additional testing on other leukemia cell lines and primary leukemia samples would help to show the range of impact in the disease. While only the top three RPs shifted cells to the γH2AX high, RP island, there were many other compounds worth investigating further. From a clinical standpoint, the RRs **CMLD011890** and **CMLD011891** also had among the highest leukemia-specific induction of γH2AX. Additionally, the RR **CMLD011880**/**CMLD010508** (**SDS-1-021**) and the ADRs **CMLD012565** and **CMLD012600** had distinct bioactivities within their subclass; the structural determinants of these are yet unknown and could also potentially be explained by engagement with specific targets.

The subset of rocaglate pyrimidinones (RPs) with exceptional, leukemia-specific γH2AX activation identified through successive testing of 600 diversity compounds followed by 37 select rocaglates demonstrate the value of an expanded biological readout space for deconvolving the impact of small chemical changes. We anticipate this combined set of experimental and computational approaches to be adaptable for testing other promising families of structurally related molecules across a range of functional readouts, cell types, and disease states.

## Methods

### Screening Compounds

Initial screens were performed on a **<u>Diversity Set</u>** of 600 compounds selected from the BU-CMD’s in-house small molecule screening collection that was comprised of 4,445 compounds at the time of **<u>Diversity Set</u>** assembly. The **<u>Diversity Set</u>** was curated to contain a roughly proportional count of members from each of the 120 distinct structural chemotypes present in the parent screening collection. **<u>Diversity Set</u>** representatives for each chemotype (range: 1-23 compounds per chemotype; average: 5.4 members per chemotype) were randomly chosen from a preselected field of available compounds meeting a specific stock volume threshold, in order to ensure availability of follow-up material for validation and secondary assays.

Follow-up assays were performed on a hand-chosen cohort of 37 rocaglates (**<u>Supplementary Figure 3</u>**), selected to include a diverse representation of bioactive rocaglate subclasses, with additional consideration of known bioactivity, structural diversity, and available stock quantity.

### PBMC Collection and Preservation

Peripheral blood mononuclear cells (PBMCs) were obtained in accordance with the Declaration of Helsinki following protocols approved by Vanderbilt University Medical Center IRB. PBMCs were collected, isolated, and cryopreserved from approximately 20 mL of freshly drawn blood as previously described ^76^. Briefly, peripheral blood was drawn into sodium heparin anticoagulant, and PBMCs were isolated by centrifugation after layering on top of a Ficoll-Paque PLUS (GE Healthcare Bio-Sciences) gradient.

### MV411 Cell Culture

MV411 cells were obtained from ATCC (CRL-9591) and confirmed to be mycoplasma negative. MV411 was cultured in IMDM (Gibco 12440-053), supplemented to a final concentration of 10% fetal bovine serum. Cells were incubated in a water-jacketed 5% CO_2_ incubator at 37 °C and maintained at densities between 100 thousand and 1 million cells per ml of culture media, fed every other day, and passaged every 4 days.

### Fluorescent Cell Barcoding (FCB) Assays

The general protocol followed Schares et. al. Barcoding plates were prepared as described in Section 6.1 of Schares et. al. ^13^.

**<u>Diversity Set</u>** Experiment

For performing **<u>Diversity Set</u>** testing, the compound plate was prepared as follows: control compounds (3 replicates of DMSO, staurosporine, etoposide, aphidicolin, rapamycin, and nocodazole) were prepared to final concentrations listed in **<u>Supplementary Table 2</u>** and plated at 1 µL into columns 6 and 12 of 8 96 well tissue culture plates (Fisher). The **<u>Diversity Set</u>** compounds were acquired from the BU-CMD and plated at a volume of 1 µL into the remaining 80 wells of the 8 96-well tissue culture plate for a final concentration of 10 µM.

MV411 cells from suspension culture (0.5-1 million cells/mL) were acquired, pelleted, and resuspended in media to around 500,000 cells/mL at least 2 hours before start of the experiment as described in Schares et. al. Cell suspension was dispensed at 200 µL to each of the 8 96 wells plates and pipetted to mix. The plate was incubated at 37°C, 5% CO_2_ for 16 hr. The remaining steps were performed as described in Schares et. al. Briefly, cells were stained for viability with 0.04 μg/mL Ax700, fixed with 1.6% paraformaldehyde, and permeabilized with 100% ice-cold methanol. Cells were stained using eight concentrations of pacific blue and six concentrations of pacific orange (per 48 wells) for generating a unique fluorescent barcode for each well, and one concentration of AlexaFluor 750 as an internal control. Cells were pooled into one flow cytometry tube per 48-well plate (16 tubes total) for staining with the following antibody panel: c-CAS3, γH2AX, p-S6 S240/244, and p-HH3 (more information detailed in **<u>Supplementary Table 1</u>**). Compensation controls for each antibody and dye and bead controls were prepared and used for the set-up of the flow cytometer. Flow cytometry data was acquired on a 5-laser (355 nM, 405 nM, 488 nM, 561 nM, and 635 nM) BD LSR II Fortessa instrument.

**<u>Rocaglate Set</u>** Experiment

For the **<u>Rocaglate Set</u>** experiment, the compound plate was prepared as follows. Control compounds (3x DMSO, staurosporine, etoposide, **CMLD010335**, rapamycin, and nocodazole) were prepared to final concentrations listed in **<u>Supplementary Table 2</u>** and plated at 1µL in columns 6 and 12 of a 96 well tissue culture plate. The **<u>Rocaglate Set</u>** (RR = 20, ADR = 8, RP = 9) acquired from the BU-CMD was plated at a volume of 1 µL into the remaining 80 wells of the 96-well tissue culture plate at a final concentration of 10 µM.

As described above, MV411 cells from suspension culture were acquired, pelleted, and resuspended in media at least 2 hours before start of the experiment. PBMCs were thawed from cryopreservation and resuspended at ∼2 × 10^6^ cells/mL. The cell suspension was dispensed at 200µL to each of the 96 wells of 2 compound plates, one for each cell type, and pipetted to mix. The plate was incubated at 37°C, 5% CO_2_ for 16 hr. The remaining steps were performed as described above, pooling cells into two total flow cytometry tubes for antibody staining: one for PBMC and one for MV411^13^. Antibody staining panel included p-STAT3, p-STAT5, p-ERK, p-HH3, p-4EBP1, p-S6 S240/244, p-S6 S235/236, Ki67, and γH2AX (**<u>Supplementary Table 1</u>**). Flow cytometry controls were performed as above. Flow cytometry data was acquired on a four laser (405 nM, 488 nM, 561 nM, and 640 nM) Cytek Biosciences Aurora spectral flow cytometer following spectral unmixing with compensation controls.

#### Dose-Response

**CMLD012390** and **CMLD013342** were prepared at the following doses and plated at a volume of 1 µL into the first two columns of rows A-H of a 96-well plate, respectively: 0.01, 0.05, 0.1, 0.5, 1, 5, 10, and 0 µM.

As described above, MV411 cells from suspension culture were acquired, pelleted, and resuspended in media at least 2 hours before start of the experiment. The cell suspension was dispensed at 200µL to each of the two columns of the 96 well plate and pipetted to mix. The plate was incubated at 37°C, 5% CO_2_ for 16 hr. Cells were stained for viability, fixed, and permeabilized. Cells for each compound were then stained using eight concentrations of pacific blue, one concentration of pacific orange, and one concentration of AlexaFluor 750. Cells for each compound were pooled into their own flow cytometry tube for staining with the same antibody panel as the **<u>Rocaglate Set</u>** experiment (**<u>Supplementary Table 1</u>**). Flow cytometry controls and data collection were performed as in the **<u>Rocaglate Set</u>** experiment.

#### Time Course

**CMLD012390** and **CMLD013342** were prepared to a final concentration of 10 µM. As described above, MV411 cells from suspension culture were acquired, pelleted, and resuspended in media. Cell suspension was dispensed at 200 µL to the first three columns of rows A-C of a 96-well plate. A 16-hour reverse time course was begun by pipetting 1 µL of DMSO, **CMLD012390**, and **CMLD013342** into columns 1, 2, and 3, respectively of row A and pipetted to mix. After 12 h, 1 µL of DMSO, **CMLD012390**, and **CMLD013342** was dispensed into row B for the 4-h time point. Lastly, after 15 h, 1 µL of DMSO, **CMLD012390**, and **CMLD013342** was dispensed into row C for the 1-h time point. At the completion of the reverse time course, all three rows were stained for viability, fixed, permeabilized, and stained using three concentrations of pacific blue, three concentrations of pacific orange, and one concentration of AlexaFluor 750. Cells were pooled into one flow cytometry tube and stained with the following antibody panel: LC3, p-EIF2a, p-MLKL, c-CAS3, and γH2AX (*cf.* **<u>Supplementary Table 1</u>**). Flow cytometry controls were performed as above. Flow cytometry data was acquired on a three-laser (405 nM, 488 nM, and 640 nM) Cytek Biosciences Northern Lights spectral flow cytometer following spectral unmixing with compensation controls.

### FCB Data Preprocessing and Analysis

Data was uploaded and stored in Cytobank for scaling, quality control gating, compensation, and analysis of unmixed cytometry data (FCS file format). Raw median fluorescence intensity values were transformed to a hyperbolic arcsine scale. For the **<u>Diversity Set</u>** test, a cofactor of 150 was selected for all functional readouts. For the **<u>Rocaglate Set</u>** experiment, the default cofactor of 6000 was selected for all markers except for Ki67 (25000), p-S6 S240/244 (12000), p-LCK (12000), p-STAT3 (12000), p-STAT5 (25000), p-S6 S235/236 (12000), p-HH3 (12000), and p-4EBP1 (12000). For the dose-response and time course experiments, a cofactor of 6000 was selected for all functional readouts. Quality control gating (QC) consisted of QC1: FSC-A vs. FSC-H for singlets, QC2: FSC-A vs. SSC-A for intact cell bodies, QC3: FSC-A vs. Ax750 for barcode uptake control, and QC4: Ax750 vs. Ax700 for viable cells. Scaled and gated samples were compensated and then computationally deconvoluted using the DebarcodeR algorithm. The resulting FCS files for each well were uploaded to Cytobank for storage and further analysis.

### T-REX Analyses

T-REX takes a pair of dimensionally reduced maps of equal size as inputs, creates a plot that highlights hotspots of cells in phenotypic regions that are the most different between the two files, and provides a T-REX degree of difference value for each analysis performed. The dimensionality reduction tool used here was t-SNE which was performed as follows: after debarcoding, the FCS files corresponding to stained cellular events from all 48 wells for both cell types were uploaded to one Cytobank experiment and fed into a t-SNE-CUDA analysis through Cytobank (settings: channels = 11 functional readouts included in the panel, iterations = 10,000, perplexity = 60, automatic learning rate, early exaggeration = 12). FCS files containing protein measurements for all readouts with the t-SNE axes appended were exported into R for subsequent analysis. Each dimensionally reduced map was sampled such that each well and rocaglate structural subclass, when applicable, was equally represented before applying T-REX. The T-REX algorithm was applied in R using a k value of 60. A modular data analysis workflow including UMAP as the dimensionality reduction tool, K-Nearest Neighbors (KNN), and Marker Enrichment Modeling (MEM) was developed in R and is available online (https://github.com/cytolab/T-REX).

### MEM Protein Enrichment Analyses

Marker Enrichment Modeling from the MEM package (https://github.com/cytolab/mem) was used to characterize feature enrichment in the RP specific island identified. MEM normally requires a comparison of a population against a reference control, such as a common reference sample, all other cells, or induced pluripotent stem cells ^28,77,78^. Here, a statistical reference point intended as a statistical null hypothesis was used as the MEM reference. For this statistical null MEM reference, the magnitude was zero and the IQR was the median IQR of all features chosen for the MEM analysis. Values were mapped from 0 enrichment to a maximum of +10 relative enrichment. The contribution of IQR was zeroed out for populations with a magnitude of 0.

### Cell Gating Analyses

Gating was performed in Cytobank to quantify the % γH2AX positive cells in response to each compound in MV411 and PBMC. This analysis was performed by plotting Side Scatter Area vs. γH2AX for each compound and cell type and using the polygon gate tool to draw an area that maximizes the difference in percentage % γH2AX positive cells between the vehicle and control wells. Quantifying the percentage of cells in the RP island was also performed using the polygon gate in Cytobank.

### EC_50_ Calculations

EC_50_s were calculated in R using the drm function from the drc package. A four-parameter log-logistic function was selected with the following unfixed parameters: hill slope, minimum value, maximum value, and EC_50_. The dependent variable for the dose-response formula was the percentage of cells in the γH2AX+ gate which was calculated on Cytobank and imported into R. The independent variable was the common logarithm of the dose for each concentration.

## Acknowledgements

Research was supported by the following funding resources: NIH/NCI grants U01 TR002526 (HLT, MJH, LEB, JAP, Jr., JMI), R01 CA226833 (JMI, HLT, MJH), HLT), R35 GM118173 (JAP, Jr.), the Vanderbilt-Ingram Cancer Center (VICC, P30 CA68485), the Michael David Greene Brain Cancer Fund (JMI), the Southeastern Brain Tumor Foundation (JMI), and the Ben & Catherine Ivy Foundation (JMI).

## Author Contributions

All authors contributed to developing the study and reviewing and editing the manuscript. LEB and JAP, Jr. developed the process to select molecules for the **Diversity Set** and **Rocaglate Set**, provided access to the **Diversity Set** and **Rocaglate Set**, and contributed to rocaglate subtype binning, R-group decomposition, and structure-activity analysis. MJH and HLT collected and processed data. HLT and JMI conducted flow cytometry data analysis and interpretation. HLT developed data analysis scripts. HLT and JMI drafted the manuscript. JMI and JAP, Jr. provided financial support.

