## Supplemental Information for "Single Cell Profiling Distinguishes Leukemia-Selective Chemotypes"

#### Supplementary Information for Thirman et. al.

**Supplementary Table 1 – Flow cytometry reagents**

| Reagent | Detect | Clone | Fluorochrome | Manufacturer | Catalog # | Readout | Diversity Set | Roc. Set + Dose Response | Time Course |
|---|---|---|---|---|---|---|---|---|---|
| Antibody | c-CAS3 | C96-605 | PE | BD Biosciences | 550822 | Apoptosis [1] | X | | X |
| Antibody | Ki67 | B56 | BV786 | BD Biosciences | 563756 | Cell proliferation [2,3] | | X | |
| Antibody | p-S6 S235/236 | D57.2.2E | Ax594 (Diversity)/Ax647 (37 Roc.) | Cell Signaling Technology | 4851S (Diversity)/9865S(37 Roc.) | mTOR+MAPK /ERK activation [4 5] | X | X | |
| Antibody | p-S6 S240/244 | D68F8 | Ax488 | Cell Signaling Technology | 5018S | mTOR activation [4 5] | | X | |
| Antibody | p-AKT S473 | D9E | APC | Cell Signaling Technology | 11962S | Upstream of mTOR [6,7] | | | |
| Antibody | p-LCK Y505 | SRRCHA | PerCP-eFluor 710 | ThermoFisher Scientific | 46-9076-42 | Src-family kinase [8] | | | |
| Antibody | γH2AX S139 | N1-431 | PerCP-Cy5.5 | BD Biosciences | 564718 | DNA damage [9] | X | X | X |
| Antibody | p-STAT3 S727 | 49/p-Stat3 | PE | BD Biosciences | 558557 | Transcriptional Activity [10] | | X | |
| Antibody | p-STAT5 Y694 | SRBCZX | PE-eFluor610 | Invitrogen | 61-9010-42 | Cell survival [11 12] | | X | |
| Antibody | p-ERK T202/Y204 | 6B8B69 | PE-Cy5 | Biolegend | 369514 | MAPK activation [13,14] | | X | |
| Antibody | p-HH3 S28 | HTA28 | PE-Cy7 | Biolegend | 641011 | M-phase cell cycle [15] | X | X | |
| Antibody | p-4EBP1 T37/46 | 236B4 | Ax647 | Cell Signaling Technology | 5123S | Translation [16] | | X | |
| Antibody | p-MLKL | D6H3V | Ax568 | Cell Signaling Technology | 91689 | Necroptosis [17] | | | X |
| Antibody | LC3 | D3U4C | Ax488 | Cell Signaling Technology | 13082 | Autophagy [18] | | | X |
| Antibody | p-EIF2α | 119A11 | Ax647 | Cell Signaling Technology | 3597 | Unfolded protein response[19] | | | X |
| NHS dye | Primary amines | | Pacific Blue | ThermoFisher Scientific | P10163 | | X | X | X |
| NHS dye | Primary amines | | Pacific Orange | ThermoFisher Scientific | P30253 | | X | X | X |
| NHS dye | Primary amines | | Ax750 | ThermoFisher Scientific | A20011 | | X | X | X |
| NHS dye | Primary amines | | Ax700 | ThermoFisher Scientific | A20010 | | X | X | X |

Table lists functional protein states used for measuring specific pathways by flow cytometry, their clone, fluorochrome, vendor source, their role in various intracellular signaling pathways, and the corresponding panel where they were used. Modeled after Balsamo et. al. [19].

**Supplementary Table 2 – Positive and negative control compounds**

| Compound | Source | Catalog # | Conc. | Purpose |
|---|---|---|---|---|
| Dimethyl sulfoxide (DMSO) | Fisher | BP231-1 | | Negative control |
| staurosporine | LKT Labs | S7600 | 1 µM | c-CAS3 + γH2AX pos. control |
| etoposide | Cayman Chemicals | 33419-42-0 | 10 µM | γH2AX + c-CAS3 pos. control |
| **CMLD010335** | BU-CMD | | 10 µM | Rocaglate control |
| rapamycin | LC Labs | NC9163747 | 0.01 µM | p-S6 suppression pos. control |
| Nocodazole | Acros | 358240100 | 4 µM | G2 cell cycle arrest + p-HH3 pos. control |
| aphidicolin | Fisher | AC611970010 | 4 µM | G1 cell cycle arrest + p-HH3 pos. control |

Table lists compounds used as positive and negative controls for specific pathway measurements made by flow cytometry, vendor source, final concentration, and a short description of their purpose.

**Thirman et. al. – Supplementary Figure 1**

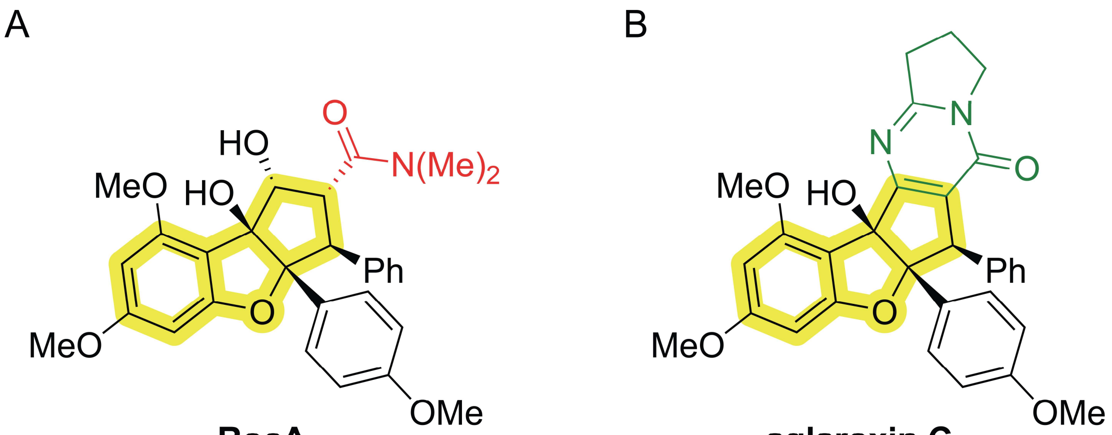


**RocA** **aglaroxin C**

**Supplementary Figure 1 – RocA and aglaroxin C are exemplar rocaglates for their respective subclasses. A)** Chemical structure for rocaglamide (RocA), a regular rocaglate (RR). R group is colored in red based on classification as RR. **B)** Chemical structure for aglaroxin C, a rocaglate pyrimidinone. Ring fusion is colored in green based on classification as RP.

#### Thirman et. al. – Supplementary Figure 2

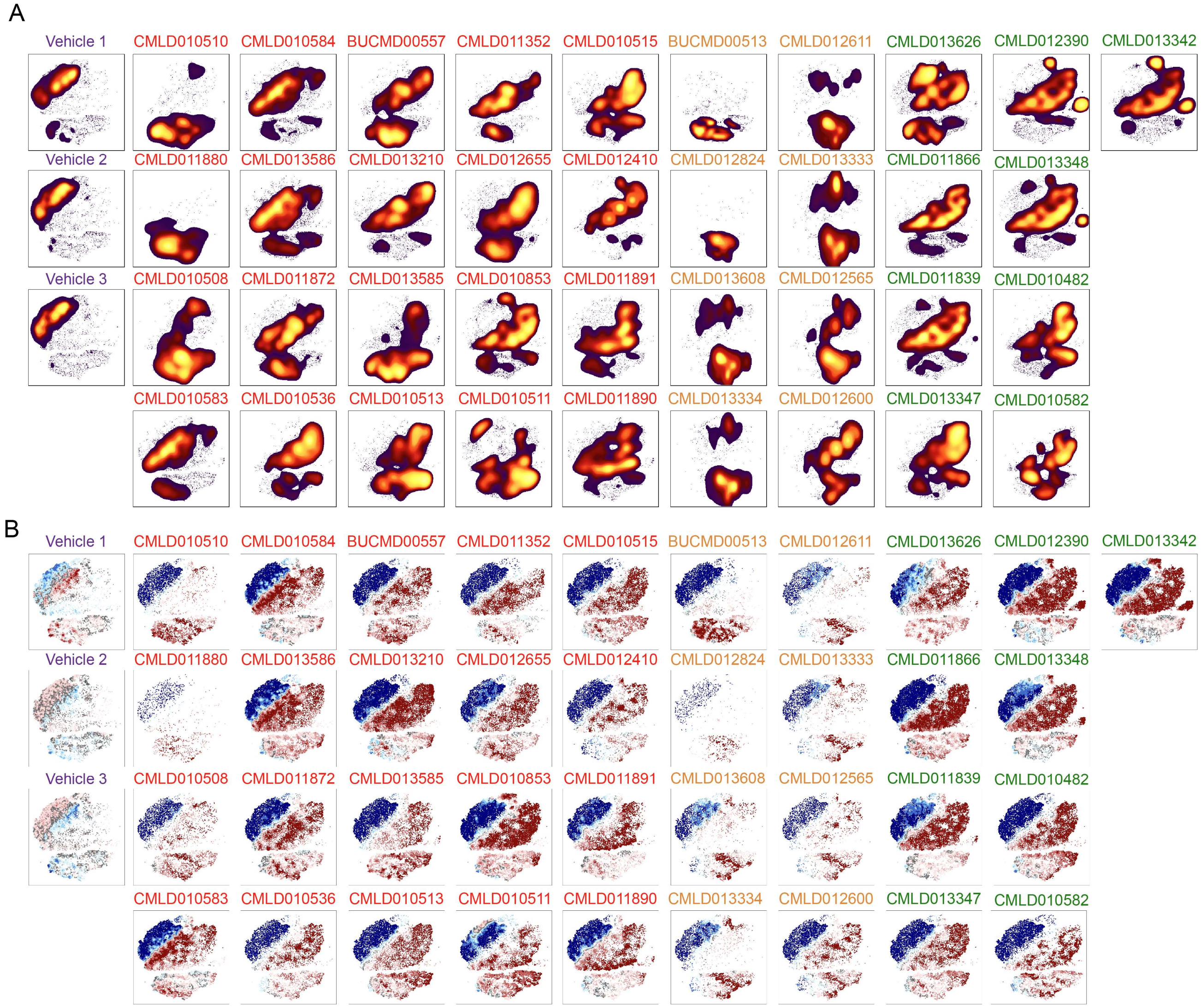


**Supplementary Figure 2 - Rocaglates had distinct patterns of bioactivity in leukemia. A)** t-SNE plots depicting the result of performing a t-SNE analysis on the entire pre-processed MV411 dataset and dividing based on compound. **B)** T-REX plots depict regions of significant difference between the t-SNE of one compound vs. the pooled set of vehicle-treated cells in MV411. For both **A)** and **B)** compound names are colored according to rocaglate structural subclass.

#### Thirman et. al. – Supplementary Figure 3

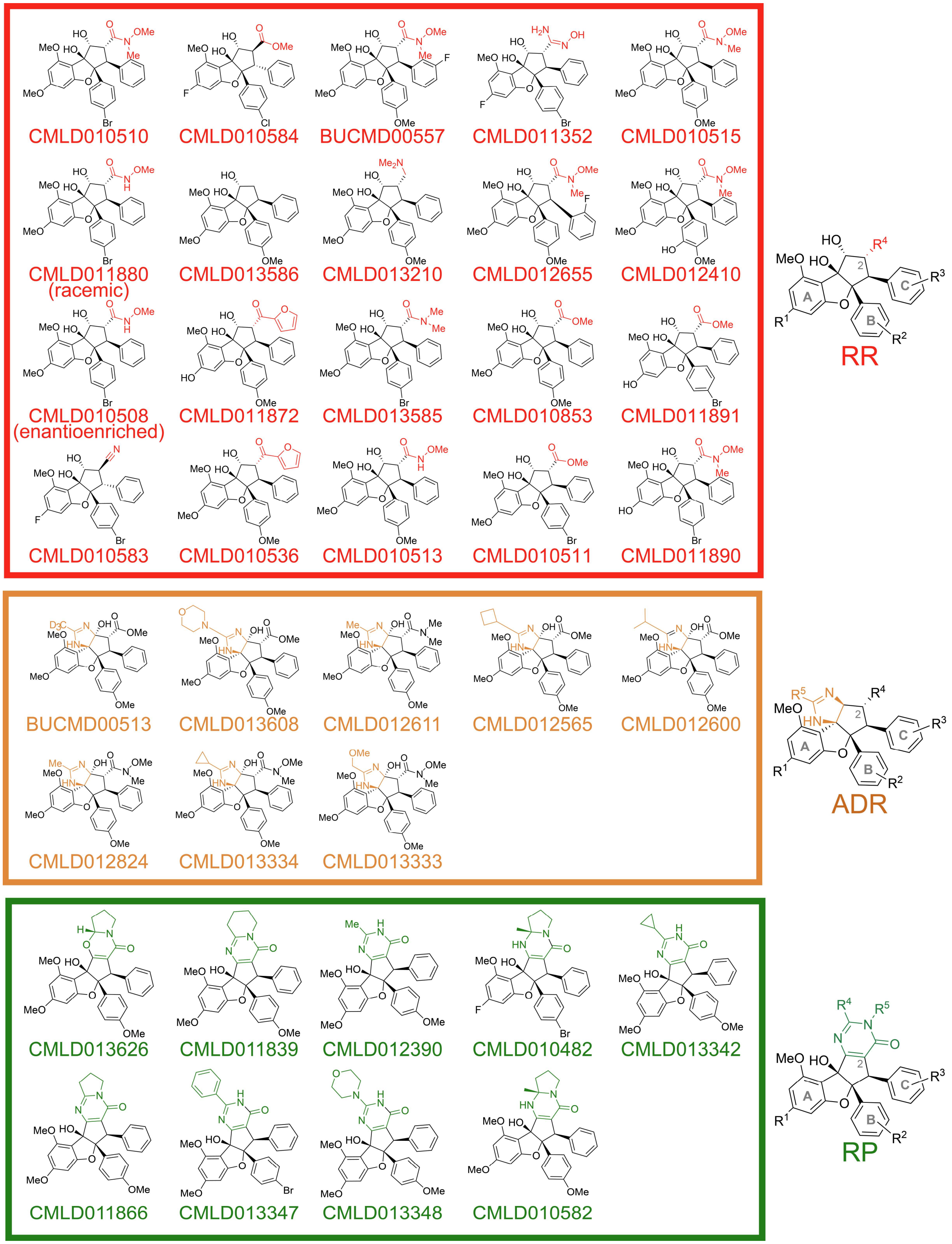


**Supplementary Figure 3 - The 37 rocaglates were organized into three structural subclasses.** The structures of the 37 rocaglates are organized into boxes according to membership to one of three subclasses. Boxes, compound names, and R groups are colored according to rocaglate subclass. The defining structural scheme for each subclass is shown to the right of each box.

**Thirman et. al. – Supplementary Figure 4**

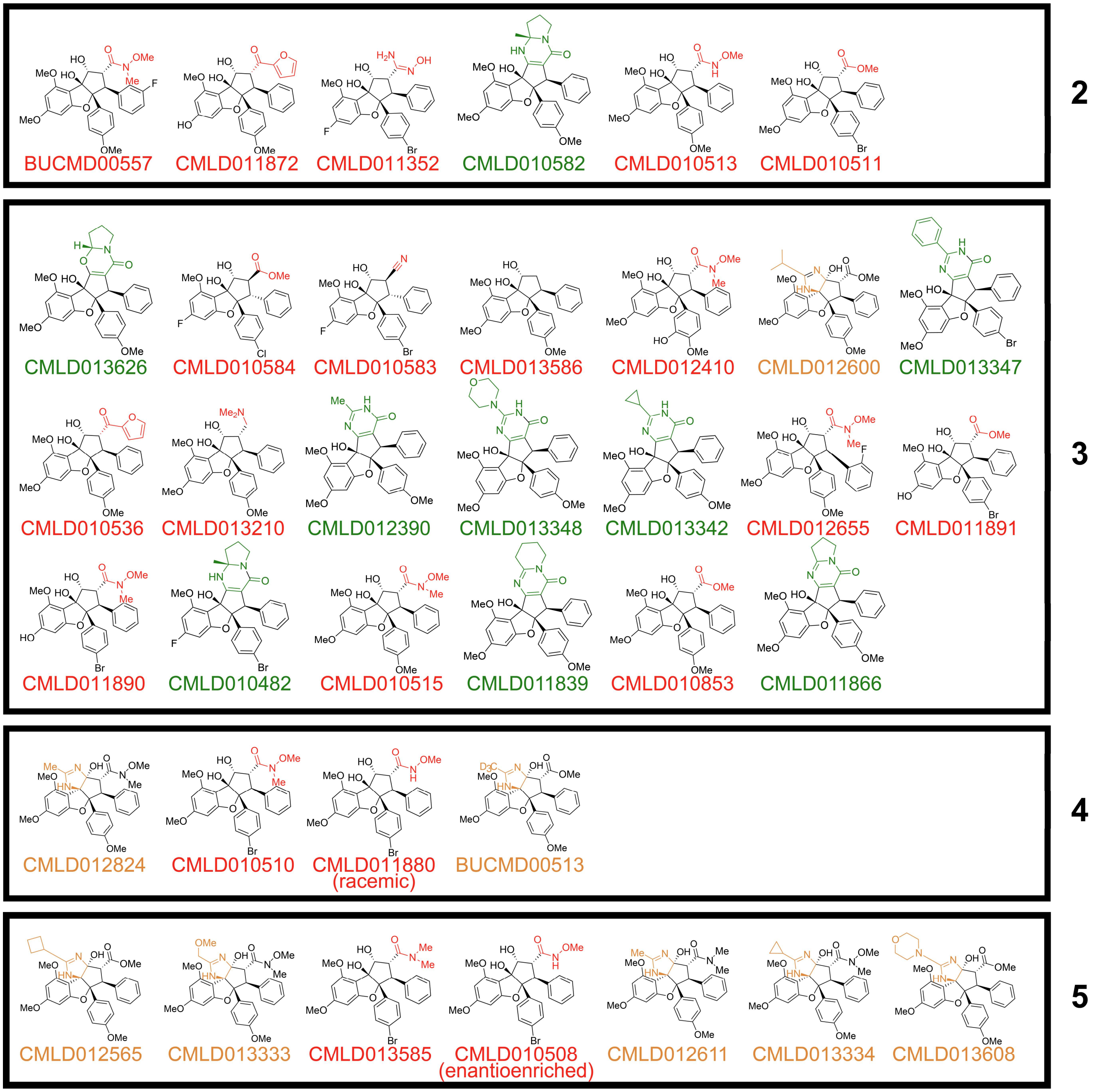


**Supplementary Figure 4 – Dendrogram clusters provided deeper insight into potential machine-learned structure-activity relationships.** The structures of the 37 rocaglates are organized according to the dendrogram clusters formed based on transformed median fluorescence intensity in **Figure 3B**. Compound names and R groups are colored according to rocaglate structural subclass. The cluster number from **Figure 3B** is listed on the right side of each box.

**Thirman et. al. – Supplementary Figure 5**

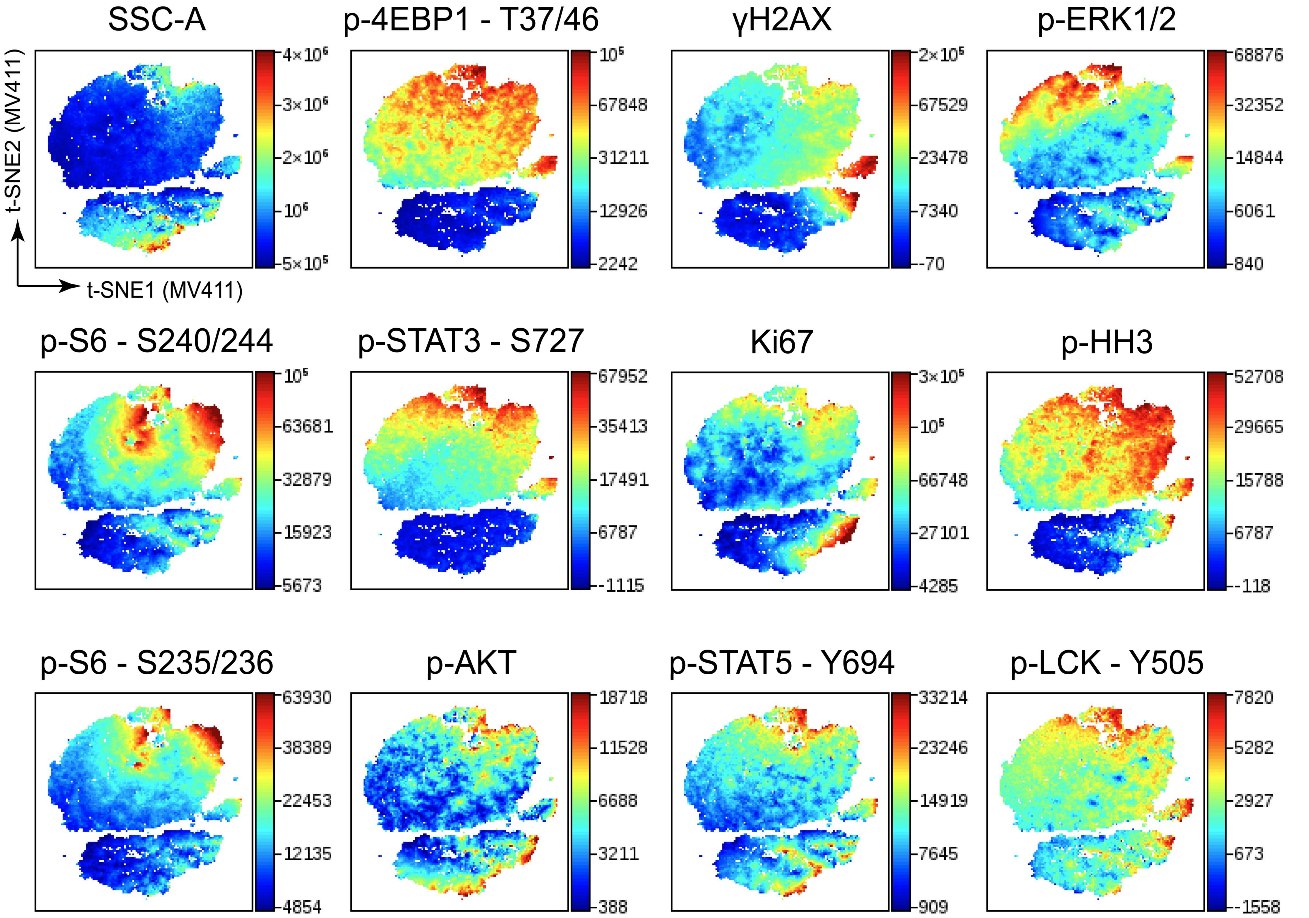


**Supplementary Figure 5 - Heterogeneity existed at the single-cell level across functional readouts.** Plot depicting the result of performing a t-SNE analysis on the entire pre-processed MV411 dataset and coloring based on protein measurements for each of the 11 functional readouts tested (and SSC-A). Each readout is on its own scale as seen in the legend on the right side of each plot.

### Thirman et. al. – Supplementary Figure 6

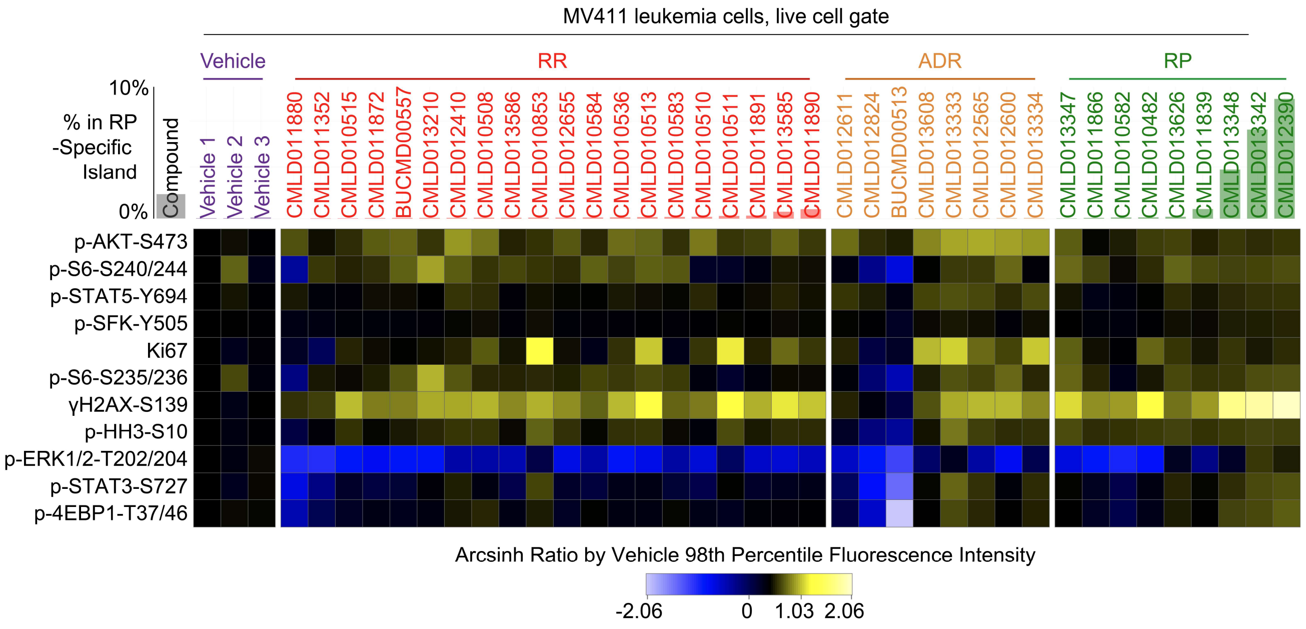


**Supplementary Figure 6 – The RP island was associated with high γH2AX and p-4EBP1 and low p-ERK.** Heatmap depicting the arcsinh ratio of the 98th percentile fluorescence intensity for each compound (listed on top of heatmap) and readout (listed left of heatmap) by the median fluorescence intensity of Vehicle 1. Cells on the heatmap range from light blue for the lowest values to bright yellow for the highest values. Compounds are grouped and colored according to the rocaglate subclass listed on the top of the plot. A bar plot depicting the percentage of cells in the RP island for each compound is shown behind each respective compound name.

**Thirman et. al. – Supplementary Figure 7**

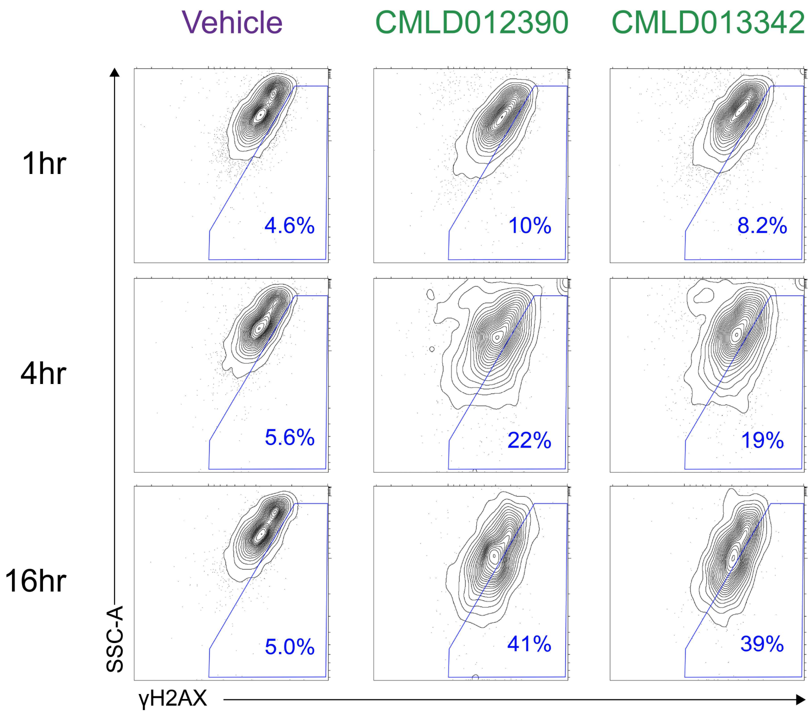


**Supplementary Figure 7 - RPs showed γH2AX activation within 4 hours that continued to increase at 16 hours.** Contour plots depicting SSC-A vs. γH2AX with 10% of cells per contour. Rows are arranged in order of increasing compound stimulation time. Percentages in the lower right corners of individual plots indicate the percentage of cells within the γH2AX+ polygon gate shown in blue. Compound names at the top of plots are colored based on membership to RP subclass.
